# A Deep Phylogenetic Split Between Short- and Long-Wavelength-Tuned Visual Systems in Surgeonfishes (Acanthuridae)

**DOI:** 10.64898/2026.11.02.763787

**Authors:** Abigail Shaughnessy, Valerio Tettamanti, Martin Luehrmann, Fabio Cortesi

## Abstract

The light environment, coupled with ecology, is widely thought to drive visual adaptation, creating species-specific visual niches. However, coral reef ecosystems experience continuous spatiotemporal variations in light conditions, and mechanisms such as phenotypic plasticity may be essential for coral reef animals to adjust to such changing sensory requirements. It remains unclear how widespread colour vision plasticity is, or the extent to which ecology and phylogeny drive plasticity in coral reef fishes. Using a comparative transcriptomic approach across 22 species, we aimed to investigate visual plasticity within a prominent coral reef fish family, surgeonfishes (Acanthuridae), in response to changes in the light environment throughout development and across seasons in adults. Surprisingly, we uncovered a deep phylogenetic split (∼42 mya) in visual opsin gene expression profiles, with Nasinae expressing a shorter-shifted palette than Acanthurinae, which coincides with the divide between water-column and benthic-feeding modalities during the speciation of their last common ancestor. However, opsin gene expression showed little variation in response to seasonal or ontogenetic shifts in the light environment, nor in the broad functional feeding groups extant species from both subfamilies fall under. Thus, the invariable visual gene expression observed in present-day surgeonfishes appears to be dictated by phylogenetic inertia, despite rapid diversification into a range of ecological niches. This suggests that spectral tuning into two main modalities, short and long, reduces the need for visual plasticity, allowing larger and more mobile coral reef fishes to conduct visually guided behaviours across the entire range of light environments they encounter.

## Introduction

Evolutionary adaptations have enabled species to partition ecological niches, reducing competition and allowing coexistence (Schoener, 1974). Such specialisations to minimise niche overlap have occurred, amongst others, at the morphological [e.g., feeding strategies in cichlids (Albertson et al., 2005)], behavioural [e.g., activity timing in lizards (Reyes-Puig et al., 2025)], and physiological [e.g., thermal tolerance in hummingbirds (Projecto-Garcia et al., 2013)] levels. Notably, underlying this phenotypic disparity are molecular differences in both gene sequence and regulation (Romero et al., 2012). Sensory system diversification, such as in olfaction, hearing or vision (Oteiza & Baldwin, 2021), is a prominent example of how trait variation can contribute to niche partitioning. Vision is a primary interface through which many animals interact with their environment and maintain optimal behavioural performance, including navigation, communication, and foraging (Cronin et al., 2014). According to the sensory drive hypothesis (Endler, 1992), fine-tuning of visual systems allows sympatric species to exploit the same ecological niche by optimising their vision to species-specific ecological demands. While direct evidence for sensory drive is exceedingly difficult to find (Cummings & Endler, 2018; Fuller & Endler, 2018), prominent examples include the long-wavelength-shifted colour vision and red nuptial male colour patterns of sticklebacks [(Brock et al., 2018); albeit, stickleback food and their light environment are also red-shifted (Bakker et al., 2024; Rennison et al., 2016)], honey bee colour vision tuned for efficient flower foraging [(Raine & Chittka), 2007; though flowering plants evolved after the bee ancestor acquired colour vision], and habitat-associated divergence of visual sensitivity in Lake Victoria cichlids [(Carleton & Kocher, 2001; Maan et al., 2006); although current-day light habitats do not reflect the light environments when cichlid colour vision first started diverging (Carleton et al., 2016; Spady et al., 2005)]. This suggests that a given animal’s visual system serves both general-purpose vision and the optimisation of specific tasks (Carleton et al., 2020; Marshall et al., 2015). However, we often lack phylogenetic context or detailed ecological information, making it difficult to assess how both factors interact to shape animal vision.

Coral reef environments are both structurally complex and rich in ecological interactions. Thus, space and resources are highly contested, providing the conditions for niche partitioning at small spatial scales (Connolly et al., 2005). At first glance, coral reef fishes share many fundamental similarities in their visual systems (Cortesi et al., 2020). For instance, visual pigments play an essential role in determining the spectral sensitivity of photoreceptors in the retina. The pigment’s wavelength of maximum absorbance (λmax) is determined by the interaction between a vitamin A-derived light-absorbing chromophore (Toyama et al., 2008; Yokoyama & Yokoyama, 1996), which is covalently bound to the opsin, a G-coupled transmembrane protein (Hunt & Collin, 2014). Most fishes have two kinds of photoreceptor cells, rods for dim light vision and cones for bright light and colour vision, which house five ancestral types of opsin proteins. Whilst rods only express the RH1 opsin [rhodopsin, λmax = 447-525 nm, PR0 (PR for photoreceptor)], cones can express SWS1 (short-wavelength sensitive 1, λmax = 347-383 nm, PR4), SWS2 (short-wavelength sensitive 2, λmax = 397-482 nm, PR3), RH2 (medium wavelength-sensitive λmax = 453-537 nm, PR2), and LWS (long-wavelength sensitive, λmax = 501-573 nm, PR1) (reviewed in Baden et al., 2025 and Carleton et al., 2020). In teleost evolution, opsin gene duplications, deletions, mutations, and gene conversion have resulted in the diversification of both opsin sequences and expression patterns (reviewed in Musilova et al., 2021). Environmental and behavioural factors, such as habitat complexity (Luehrmann et al., 2020), feeding ecology (Stieb et al., 2023, 2024), and colour patterns (Tettamanti et al., 2025), have been proposed as evolutionary drivers for the diversity in opsin gene sequence and expression observed in coral reef fishes today.

Importantly, the underwater light environment is not static. The spectral characteristics of the habitat can differ with depth (Jerlov, 1977; Loew & McFarland, 1990), diurnal cycles, seasonality (Loew & McFarland, 1990), and on an even smaller microhabitat-scale (Luehrmann et al., 2020; Marshall et al., 2003b). Thus, to navigate these dynamic light environments, many fishes, but not all (Tobler et al., 2010; Valen et al., 2017), exhibit phenotypic plasticity in their visual system (Fuller & Claricoates, 2011; Luehrmann et al., 2018; Nandamuri et al., 2017; Stieb et al., 2016). Phenotypic plasticity, encompassing both developmental plasticity and phenotypic flexibility (i.e., reversible phenotypic change within an individual’s lifetime) in opsin gene expression, has been documented in several freshwater and coral reef fish species. Examples include damselfishes (Pomacentridae) inhabiting different depths (Stieb et al., 2016), killifish (Fundulidae) in clear versus murky waters (Fuller et al., 2005), and development changes across the lifetime of several marine and freshwater species [e.g., brown dottyback (*Pseudochromis fuscus*; Cortesi et al., 2016, spotted unicornfish (*Naso brevirostris*; Tettamanti et al., 2019), crater lake cichlids (Cichlidae; Härer et al., 2017; Karagic et al., 2019), and rainbow trout (Salmonidae; Cheng & Flamarique, 2007; Cheng & Novales Flamarique, 2004)]. However, much of this work has been restricted to single species or to small, young evolutionary radiations, as in crater lake cichlids. Hence, the question remains whether and to what extent ecological and phylogenetic factors govern visual system plasticity in closely related fish species that share the same ecological niches (Shaughnessy & Cortesi, 2024).

Surgeonfishes (Acanthuridae) have a long evolutionary history closely associated with coral reef ecosystems. The earliest major divergence occurred in the early Eocene (∼42 Ma), separating the family into two major subfamilies: Acanthuridae (tribes: Acanthurini, Zebrasomini, and Prionurini) and Nasinae (Sorenson et al., 2013). Subsequent diversification accelerated during the late Miocene–Pliocene (∼20 Ma), giving rise to the 84 extant species that perform a range of key ecosystem functions on present-day coral reefs, including controlling algal turf biomass through bottom-feeding, preventing parasitic infections, and transporting inorganic sediments (Marshell & Mumby, 2015). Despite a deep evolutionary separation and remarkable diversity in ecology, behaviour, and colouration, surgeonfishes can be categorised into six broad functional groups based on similarities in morphological traits and feeding modes: browsers, brushers, croppers, concealed croppers, sediment-suckers, and water-column feeders (Tebbett et al., 2022). Notably, these functional groups evolved in parallel several times within and between species of Acanthurinae and Nasinae (Friedman et al., 2016; Siqueira et al., 2019; Tebbett et al., 2022), potentially driving group-specific specialisations of the visual system. Supporting this, Stieb et al. 2024 reported a long-wavelength-shifted visual system in the sediment-sucking ringtail surgeonfish (*Acanthurus blochii*), which likely assists in detecting far-red-emitting algae on the benthos. Contrarily, the spotted unicornfish (*N. brevirostris*) was found to have a short-wavelength-shifted visual system, which presumably assists with detecting UV-absorbing or reflecting zooplankton/gelatinous prey items in the water column (Stieb et al., 2024; Tettamanti et al., 2019). Notably, our previous work in damselfishes suggests that divergence towards short– and long-wavelength-shifted visual systems co-occurred with the evolution of different feeding ecologies and fish colouration (Stieb et al., 2023, 2024). However, unlike the often site-attached, territorial damselfishes, many surgeonfish species occupy dynamic niches, with shifts in habitat use, diet, and responses to abiotic conditions occurring throughout development and into adulthood (Claydon et al., 2012; Marshell et al., 2011).

The visual system of the Acanthurinae, convict surgeonfish (*Acanthurus triostegus*; Besson et al., 2020; Fogg et al., 2023) and the Nasinae, *N. brevirostris* (Tettamanti et al., 2019), is plastic during development, with opsin gene expression and morphological changes in the retina taking place during settlement and metamorphosis from the larval to juvenile stages. As adults, many species maintain large home ranges and often migrate through the water column or between reefs (Tebbett et al., 2022), thus experiencing substantial daily changes in their visual environments. Over longer timescales, seasonal changes in the light environment occur across coral reefs, where winter conditions are associated with clear water with minimal suspended or dissolved matter, resulting in a broad ‘blue’ light environment, spanning wavelengths from ultraviolet (UV) to red. During summer, algae and suspended sediments dominate, making the water murkier and more red-shifted (Jerlov, 1977; Lythgoe, 1979; McFarland, 1986). Thus, the contrast between deep phylogenetic separation, strict functional groups, and continuous changes in their light habitat makes surgeonfishes ideal for investigating whether their visual systems are the product of phylogenetic inertia, have coevolved with ecological niche specialisation, or whether plasticity in the system enables them to continuously adapt to the prevailing visual environment.

To address these three scenarios, we first used a phylogenetically comparative transcriptomic approach across 22 surgeonfish species from both subfamilies to determine whether their retinal gene expression has diverged in line with their evolutionary history or reflects their functional groups. Focusing on opsin gene expression as a proxy for colour vision, we then investigated whether surgeonfish exhibit phenotypic plasticity in response to (1) developmental processes or (2) seasonal fluctuations of light in their natural habitat as adults.

## Results

To assess visual specialisation, a phylogenetically representative set of Acanthuridae (22 of 84 species; n = 108 individuals; Fig. S1) from Australia’s Northern (Lizard Island, 14°40′S, 145°27′E) and Southern (Heron Island, 23°26′S, 151°55′E) Great Barrier Reef (GBR) was investigated (Table S1). Except for *Prionurus* spp., at least one species from each major clade and genus was sampled. We used bulk retinal transcriptome sequencing (RNA-seq) to identify the visual opsin gene repertoire and quantify expression in each species. Phylogenetic reconstruction using a reference vertebrate opsin dataset confirmed that six full-length coding sequences of opsin genes are present across the surgeonfish family: *rh1, sws2b, sws2Aa, rh2c, rh2a* and *lws* (Figs. 1 and 2; Table S2 and S3). In addition, four species were found to possess *sws2b* pseudogenes: the blue tang (*Paracanthurus hepatus*) has two, and the striated surgeonfish (*Ctenochaetus striatus*), the twospot bristeltooth (*Ctenochaetus binotatus*), and the mimic surgeonfish (*Acanthurus pyroferus*) each have one. These pseudogenes contain premature stop codons at various positions in the amino acid sequence (Fig. 1).

**Figure 1:**
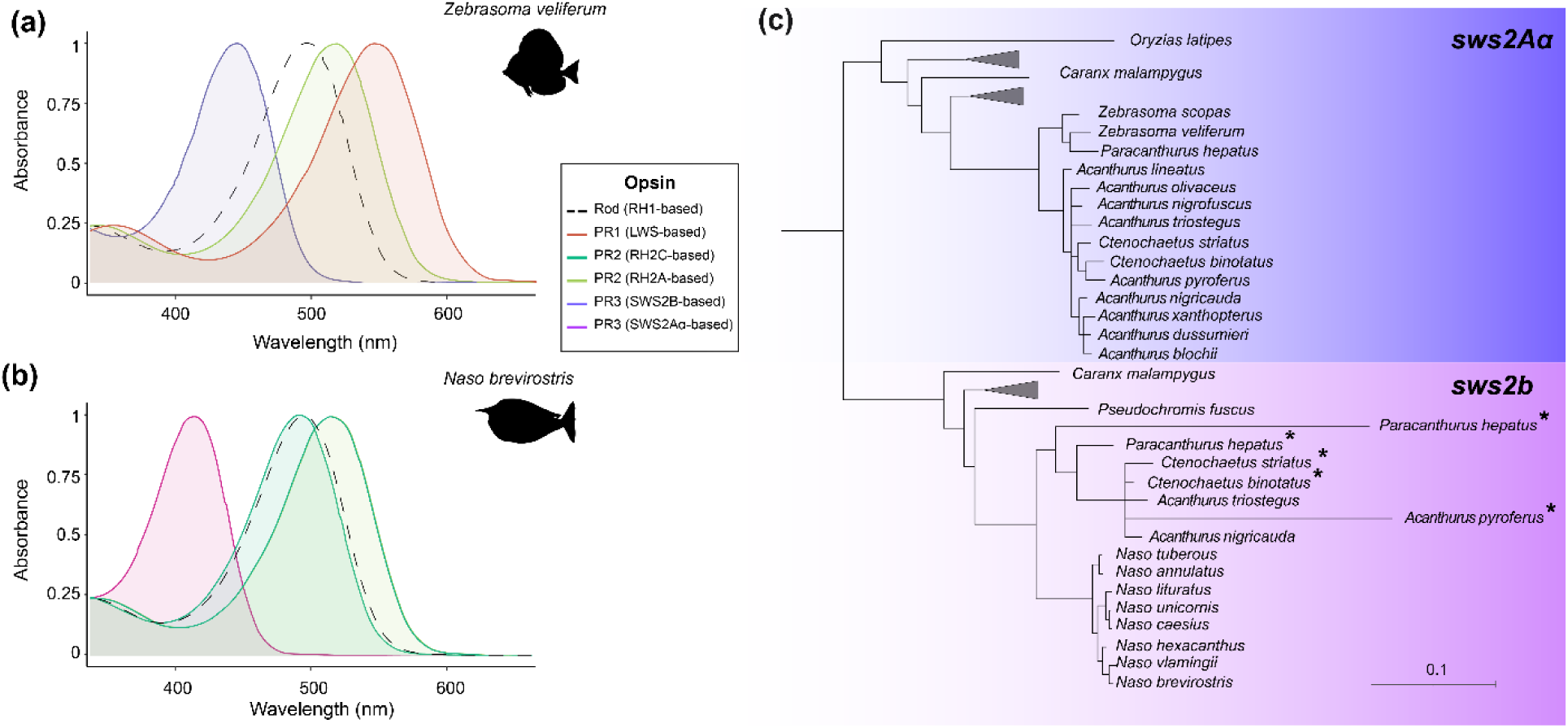
Estimated photoreceptor spectral sensitivities and short-wavelength-sensitive opsin gene phylogeny of surgeonfishes (Acanthuridae). (a, b) The estimated visual pigment spectral absorbance curves of adult surgeonfish as predicted by VPOD (Frazer et al., 2024)). (a) The Acanthurinae sailfin tang (*Zebrasoma veliferum*) is longer wavelength-shifted, using three main cone opsins: SWS2Aα (short-wavelength-sensitive, 445 nm λmax: purple), RH2A (green-wavelength-sensitive, 517 nm λmax: light green), and LWS (long-wavelength-sensitive, 547 nm λmax: red). (b) Conversely, the Nasinae spotted unicornfish (*Naso brevirostris*) is shorter wavelength-shifted, mainly using the three cone opsins: SWS2B (414 nm λmax: pink), RH2C (492 nm λmax: green), and RH2A (519 nm λmax: light green). Rods are represented by a black dashed line; peak sensitivity occurs at 496 nm in both *Z. veliferum* and *N. brevirostris*. Genes with low expression levels in both species (< 7%) are not shown (refer to Table S3 for details). (c) Short-wavelength-sensitive (*sws*) visual opsin gene phylogeny based on the retinal transcriptomes of 22 surgeonfish species. Note that Acanthurinae mostly express the longer wavelength-shifted *sws2Aa opsin,* while the Nasinae exclusively use the shorter shifted *sws2b* opsin. The ultraviolet sensitive *sws1* gene was not expressed in any surgeonfish species. *Sws2b* pseudogenes (as indicated by an *) were identified in four Acanthurinae species. The scale bar denotes substitutions per site.

**Figure 2:**
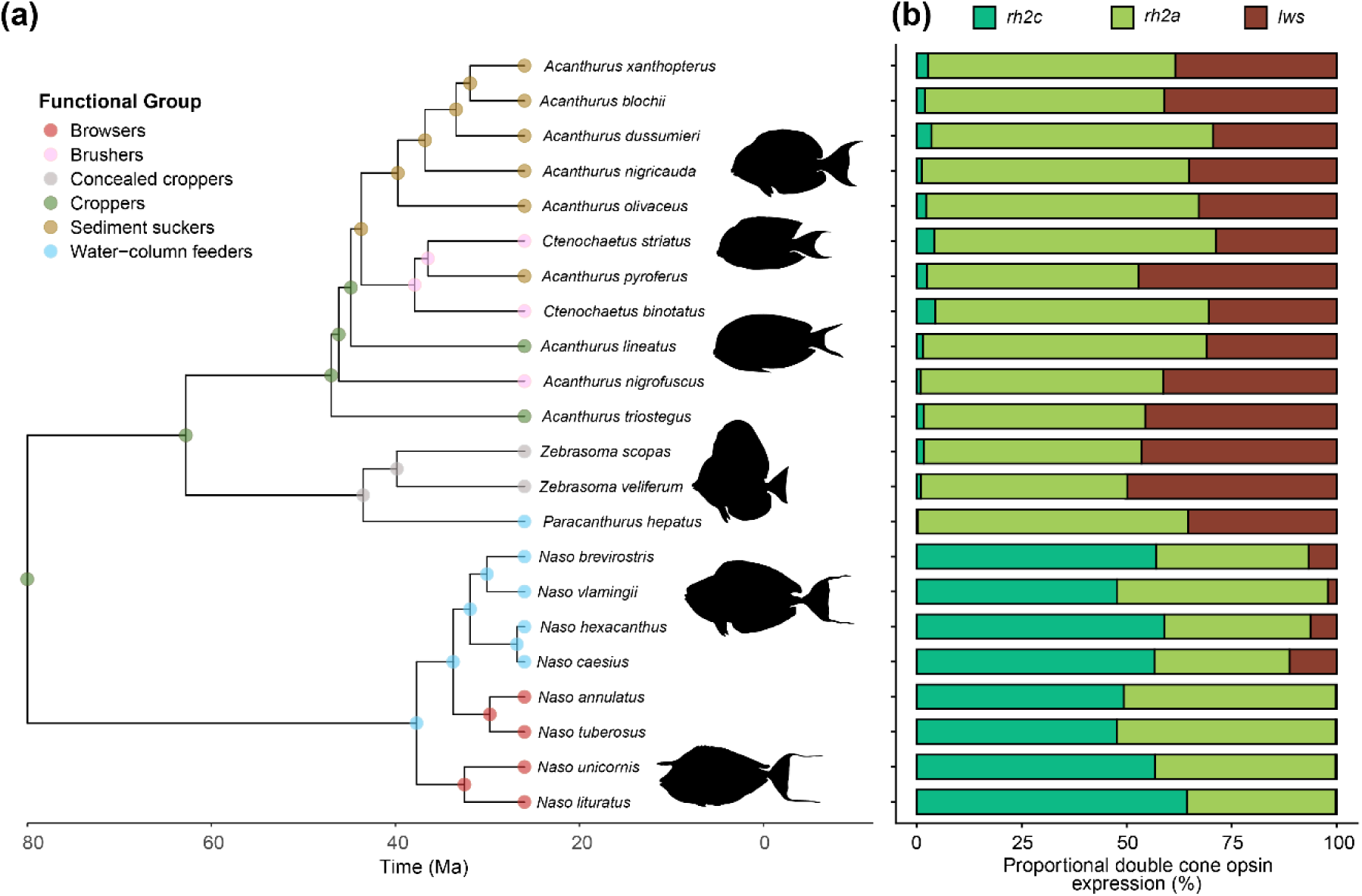
Surgeonfish species phylogeny and associated double cone opsin gene expression. (a) Phylogenetic tree of the Acanthuridae (surgeonfish) family investigated in this study (n = 22, obtained from Fish Tree of Life, Rabosky et al., 2018). Functional groups, as per Tebbett et al. (2022), are mapped to extant species and node probabilities from ancestral-state reconstruction. (b) The mean proportional expression of long-wavelength sensitive opsin genes expressed in the double cones (for details, see Table S3). Opsin expression profiles were averaged across the seasons. However, given the seasonal variation seen in sailfin tang (*Z. veliferum*), only the autumn and winter opsin expression profiles were included. Species typically expressed *rh2c* (green-wavelength sensitive, PR2), *rh2a* (green-sensitive, PR2) and *lws* (red-sensitive, PR1), with a clear ancestral split between the Acanthurinae (longer-shifted) and the Nasinae (shorter-shifted) in gene expression profiles.

### Diversity of proportional opsin gene expression across the surgeonfish phylogeny

Irrespective of what season they were sampled in, Acanthurinae expressed a longer-shifted visual opsin gene complement compared to Nasinae (Fig. 2, Table S3). *Acanthurus, Ctenochaetus,* and *Zebrasoma* species expressed the *sws2Aα* single-cone gene, with trace levels of *sws2b* expression (∼0.1% of total single cone gene expression) detected in *A. nigricauda* and *A. triostegus*. For the double-cone genes, Acanthurinae expressed high levels of *rh2a* and *lws* (Fig. 2, Table S3). All water-column feeder and browser species within the *Naso* genus highly expressed *rh2c* and *rh2a* double cone genes, and exclusively *sws2b* in the single cones. Moreover, most Nasinae species exhibited low *lws* expression. However, despite being regarded as a water-column feeder, blue tang (*P. hepatus*) showed low expression of *rh2c* in its double cones (0.3 ± 0.2%). Instead, this species demonstrated high expression of *rh2a* (64.3 ± 5.3%) and *lws* (35.4 ± 5.1%) (Fig. 2, Table S3).

### Evolutionary relationship between opsin expression and ecological traits

The single-cone gene expression pattern suggests a deep evolutionary split independent of current-day ecology, with all Nasinae having a shorter-shifted opsin variant compared to the Acanthurinae (Fig 2). This was supported by a phylogenetic generalised linear model (PGLM), which showed that feeding group was not associated with *sws2b* and *sws2Aα* expression (all p > 0.99). For the double cones, however, the pattern was less obvious (Fig. 2). Hence, to investigate potential evolutionary links between gene expression and either functional group or feeding strategy, we conducted a phylogenetic generalised least squares regression (PGLS). Strong phylogenetic dependence, but no significant relationship was observed between the expression of *rh2c* (λ = 1, F2,19 = 0.99, p = 0.39) and *lws* (λ = 1, F2,19 = 0.73, p = 0.50) in relation to the respective feeding groups. Conversely, while still showing a considerably strong phylogenetic signal, a significant association was observed between feeding category and *rh2a* expression (λ = 0.76, F2,19 = 3.74, p = 0.043).

### Spectral sensitivity (λmax) estimations of opsins

To investigate the spectral tuning of the identified opsins, we estimated the spectral sensitivities (λmax) of the visual pigments bound to an A1 chromophore across all surgeonfish species using amino-acid-based modelling (AA-modelling, as described in Cheney et al., 2022; Luehrmann et al., 2018) and a machine-learning model called the Visual Physiology Opsin Database (VPOD, Frazer et al., 2024) (Fig. 1). Estimated RH1 pigment λmax values from AA-modelling ranged from 484 nm in *C. binotatus* to 496 nm in most species (Table S4). Similarly, the VPOD estimations of the RH1 pigment ranged from 482.4 nm to 497 nm. These estimates align well with previous physiological measurements of surgeonfish rod photoreceptor spectral sensitivities using microspectrophotometry (MSP) (Losey et al., 2003), with the orange-spine unicornfish (*Naso lituratus*) exhibiting the largest difference (MSP: 497 nm, VPOD: 493.4 nm, AA-modelling: 496 nm; Table S4).

AA-modelling of the SWS2B [all species of the genus Naso); Fig. 1] single-cone visual pigment in the Nasinae predicted a maximal sensitivity at 424 nm (Table S4). In contrast, the VPOD estimates predicted the SWS2B pigment to have a λmax between 414 and 462.2 nm. These estimates closely match spectral sensitivities previously determined using MSP, where the violet cone for orange-spine unicornfish (*Naso lituratus*) and blue-spine (*N. unicornis*) measured 421 nm and 416 nm, respectively (Losey et al., 2003).

In contrast, the longer-wavelength-shifted SWS2Aα-based pigment showed greater variation between Acanthurinae species (Table S4). The estimated spectral sensitivity of SWS2Aα pigments from AA-modelling ranged from 444-448 nm in blue tang (*P. hepatus*: VPOD: 421.9 nm) to 456-464 nm in the brown tang (*Zebrasoma scopas*; VPOD: 445.9 nm), with the pigments found in the remaining species falling within this range (Table S4). MSP measurements found one type of single cone maximally sensitive at 458 nm in convict tang (*A. triostegus*: Losey et al., 2003), which agreed with the estimated λmax of SWS2Aα for this species (AA-modelling: 452-460 nm). MSP in the sailfin tang (*Zebrasoma veliferum*) found its single cones to be maximally sensitive at 448 nm (Losey et al., 2003). However, AA-modelled λmax estimation for SWS2Aα in this species was noticeably higher (456-464 nm) than MSP measurements, whereas the VPOD estimation showed a closer prediction (444.8 nm) (Table S4).

Calculated λmax of surgeonfish RH2C pigments with AA-modelling were again highly variable, ranging from 467-476 nm (lower-upper boundary) in the ringtail unicornfish (*Naso annulatus*) and the humphead unicornfish (*Naso tuberosus*) to 475-484 nm in the mimic surgeonfish (*A. pyroferus*). In contrast, the VPOD estimates of the RH2C pigment showed less variation between species, ranging from 489.7 to 492.9 nm (Table S4). Of the four species for which double and/or twin cone photoreceptor spectral sensitivities have been determined by MSP, all VPOD predictions were closer to the MSP values (Losey et al., 2003).

Predicted λmax for surgeonfish RH2A pigments showed little variability and ranged from 510-513 nm in twospot bristletooth (*C. binotatus*) to 514-516 nm in all species of the genus *Naso*, with all other species falling within this range (Table S4). Likewise, little variation was observed in the VPOD estimates for the RH2A pigment (516.5-518.6 nm). Our AA-modelled estimates mostly underestimate surgeonfish RH2A λmax compared to previous work on four species using MSP, in which one spectrally distinct double/twin cone type in *A. triostegus* (518 nm), two in *N. lituratus* (495/521 nm) and bluespine unicornfish (*N. unicornis*: 486/507 nm), and three in sailfin tang (*Z. veliferum*: 511/517/523 nm) were found (Losey et al., 2003).

Estimated λmax for surgeonfish LWS pigments ranged from 544 nm in *N. lituratus*, the bluespine unicornfish (*N. unicornis*) and the bignose unicornfish (*N. vlamingii*), over 553 nm in mimic surgeonfish (*A. pyroferus*) to 560 nm in all other tested surgeonfish species (Table S4). Compared to double/twin cone λmax in four surgeonfish species determined by MSP (Losey et al., 2003), these estimates are ∼ 20 nm longer-shifted than even the highest values determined by MSP (Table S4).

### Seasonal light environment shifts and flexibility in opsin gene expression

Next, we examined the phenotypic flexibility of opsin gene expression in response to southern-hemisphere seasonal shifts in the light environment. To this end, an additional subset of seven surgeonfish species from both subfamilies was sampled during summer (December to February), autumn (end of March to May) and winter months (July and August; 3-5 adult individuals per season; Table S1). We then measured the absolute spectral irradiance at 1-2 m depth underwater between seasons at a single reef location at Lizard Island (Fig. S2). The spectra during winter showed a small difference from those in summer, in which shorter wavelengths, including UV, were mostly attenuated in summer. In addition, the light intensity during summer was higher compared to winter. To gain further insight into year-round variation in the light environment and temperature (two key drivers of algal growth; Singh & Singh, 2015), we examined long-term sensor data collected by the Australian Institute of Marine Science (AIMS). This data demonstrated cyclic fluctuations in the underwater Photosynthetically Active Radiation (PAR 400-700 nm) and temperature, with the highest recordings occurring during summer (Fig. S2). Retinal opsin gene expression revealed that, except for sailfin tang (*Z. veliferum*), surgeonfishes showed no significant season-associated differences in the relative expression of opsin genes (Fig. 3, Table S5). Sailfin tang (*Z. veliferum*), however, increased the expression of the red-sensitive *lws* (n = 3 per season: Summer, 56.6 ± 4.53 %; Winter, 38.90 ± 1.85%; z = 2.53, p = 0.03) and decreased the expression of their green-sensitive *rh2a* opsin in summer (Summer, 43.20 ± 2.62 %; Winter, 59.10 ± 2.00 %; z = 2.53, p = 0.03). There was no significant seasonal variation in *rh2a* or *lws* expression when comparing spring to winter (Spring, 44.0 ± 2.5%; z = 1.04, p = 0.3, n = 3), and summer (z = –1.49, p = 0.3) (Fig. 3b and 3c, Table S5).

**Figure 3:**
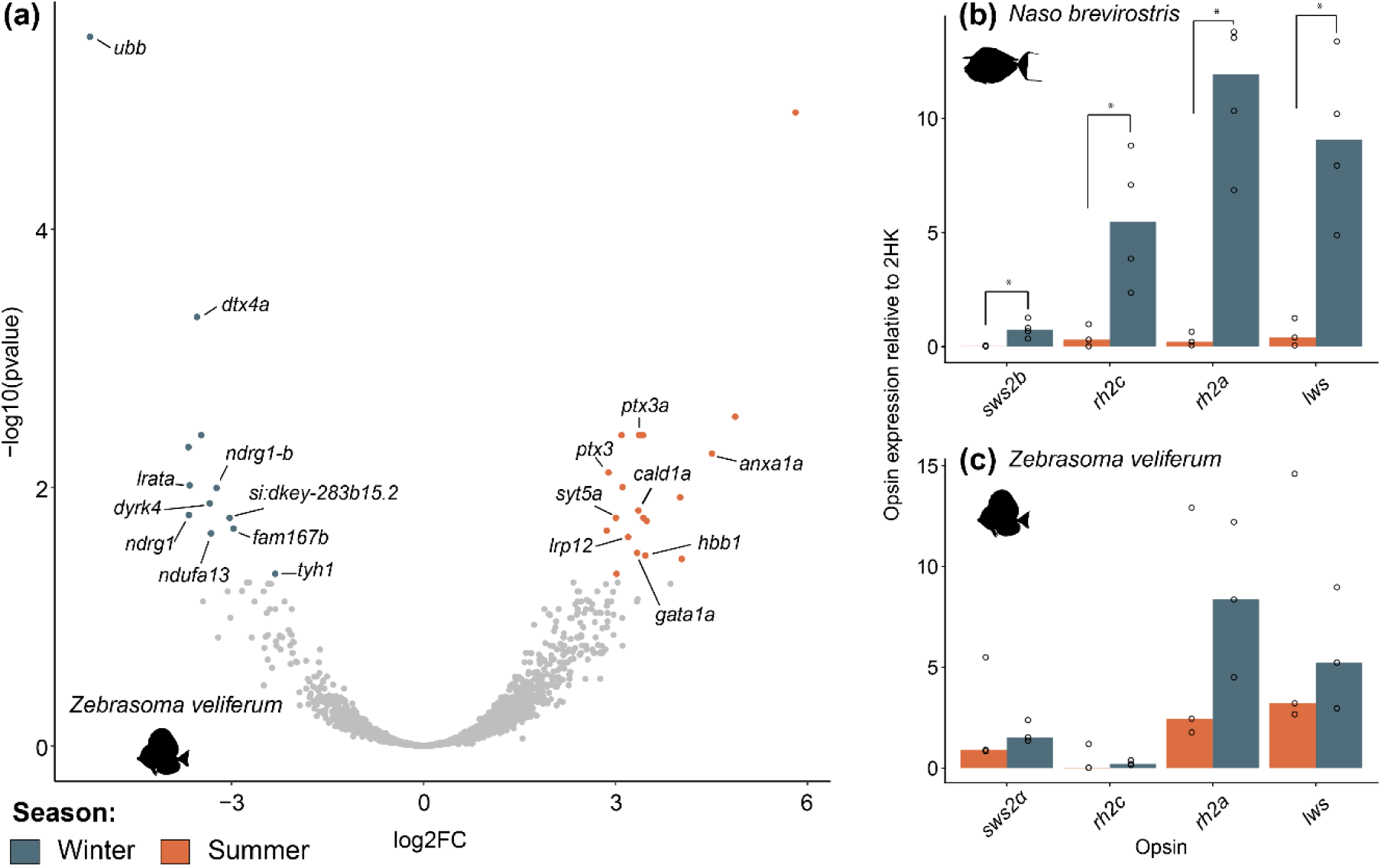
Differential retinal gene expression across seasons in surgeonfishes: (a) Volcano plot illustrating the differentially expressed retinal genes between summer and winter in sailfin tang (*Zebrasoma veliferum*). The fold changes (FC) in the expression of all genes were plotted against the (–log10) adjusted p-value. The top 15 up– and downregulated genes (i.e. greatest fold change) across the seasons are labelled (see supplementary Tables S7 and S8 for details). Orange dots with a log fold change > 0 represent genes that were expressed at significantly higher levels in summer than winter, whilst blue dots represent genes with a log fold change < 0. Grey dots are genes that showed no seasonal difference in expression. (b,c) Relative changes in opsin gene expression, normalised to two housekeeping genes [β-actin (*actb*) and elongation factor-1a (*elf1a*)], are shown for the spotted unicornfish (*Naso brevirostris*) and *Z. veliferum* (see supplementary Table 6). The height of the bars represents the median expression. Opsin gene expression was compared across seasons using a Mann-Whitney test (*p < 0.05).

We further investigated seasonal changes in relative opsin gene expression by normalising it to two stably expressed housekeeping genes, β-actin (*actb*) and elongation factor-1α (*elf1a*) (Yourick et al., 2019). Several species, including striated surgeonfish (*C. striatus*), sailfin tang (*Z. veliferum*), bluespine unicornfish (*N. unicornis*), showed a trend towards upregulation of opsin gene expression in winter (Fig. 3c). However, this trend was significant in the spotted unicornfish (*N. brevirostris*) (W= 0, p = 0.03, Fig 3b). In contrast, the lined surgeonfish (*Acanthurus lineatus*) showed a slight upregulation of *rh2a* and *lws* in summer. Finally, eyestripe surgeonfish (*A. dussumieri*) and epaulette surgeonfish (*A. nigricauda*) showed no seasonal change in opsin gene expression (Table S6).

### Seasonal differential gene expression of the whole retina

We next investigated which retinal genes, beyond the opsins, exhibited significant seasonal changes in expression. We performed a differential gene expression analyses on all retinal genes present in two surgeonfish species: sailfin tang (*Z. veliferum*), which exhibited seasonal shifts in the proportional expression of opsins, and spotted unicornfish (*N. brevirostris)*, which showed stable opsin expression and served as a non-shifting control (Fig. 3a, Table S7 and S8). The seasonal comparison found 155,847 differentially expressed transcripts in *Z. veliferum* and 103,439 transcripts in *N. brevirostris.* The subsequent pre-processing allowed 23,278 and 21,645 genes of *Z. veliferum* and *N. brevirostris* to pass the filtering, respectively. In summer, both *Z. veliferum* and *N. brevirostris* upregulated genes associated with oxygen carrier activity (*hba*, *hbb1* and *hbb2*), DNA/RNA binding (*gata1a, rpl19* and *rpl30*) and calcium ion binding (*anxa1a, anxa5* and *rpl17*). In addition, *Z. veliferum* in summer downregulated genes that are involved in transferase activity (*lrata* and *dtx4a*) and mitochondria (*ndufa13, fam167b,* and *si:dkey-283b15.2*). Conversely, for *N. brevirostris,* the genes downregulated in summer are associated with protein kinase activity (*camk1gb*), transcriptional coregulation activity (*nrip1b*), and SNAP receptor activity (*vamp2*) (Table S7 and S8). Notably, during the summer months, genes involved with outer segment photoreceptor morphology (*ndrg1* and *ndrg1-b*) were also downregulated in both species (Fig. 3a, Table S7 and S8).

### Developmental plasticity in visual opsin gene expression

Proportional expression of opsin genes at different life stages was assessed for four species: epaulette surgeonfish (*A. nigricauda*), bignose unicornfish (*N. vlamingii*), brown tang (*Z. scopas*), and sailfin tang (*Z. veliferum*)*. Naso vlamingii* expression was analysed in three stages [larval, juvenile (autumn) and adult (winter)], while only two stages were investigated in *A. nigricauda*, *Z. scopas*, and *Z. veliferum* (juveniles and adults collected in autumn). In *A. nigricauda*, *Z. scopas*, and *N. vlamingii*, the proportional expression of individual opsins showed no significant differences between life stages. In contrast, *Z. veliferum* within one autumn season showed increased *rh2a* expression in adults compared to juveniles (n = 3 per stage: Juvenile, 61.35 ± 2.09 %; Adult, 54.33 ± 4.23%; z = 9, p = 0.05) (Fig. 4).

**Figure 4:**
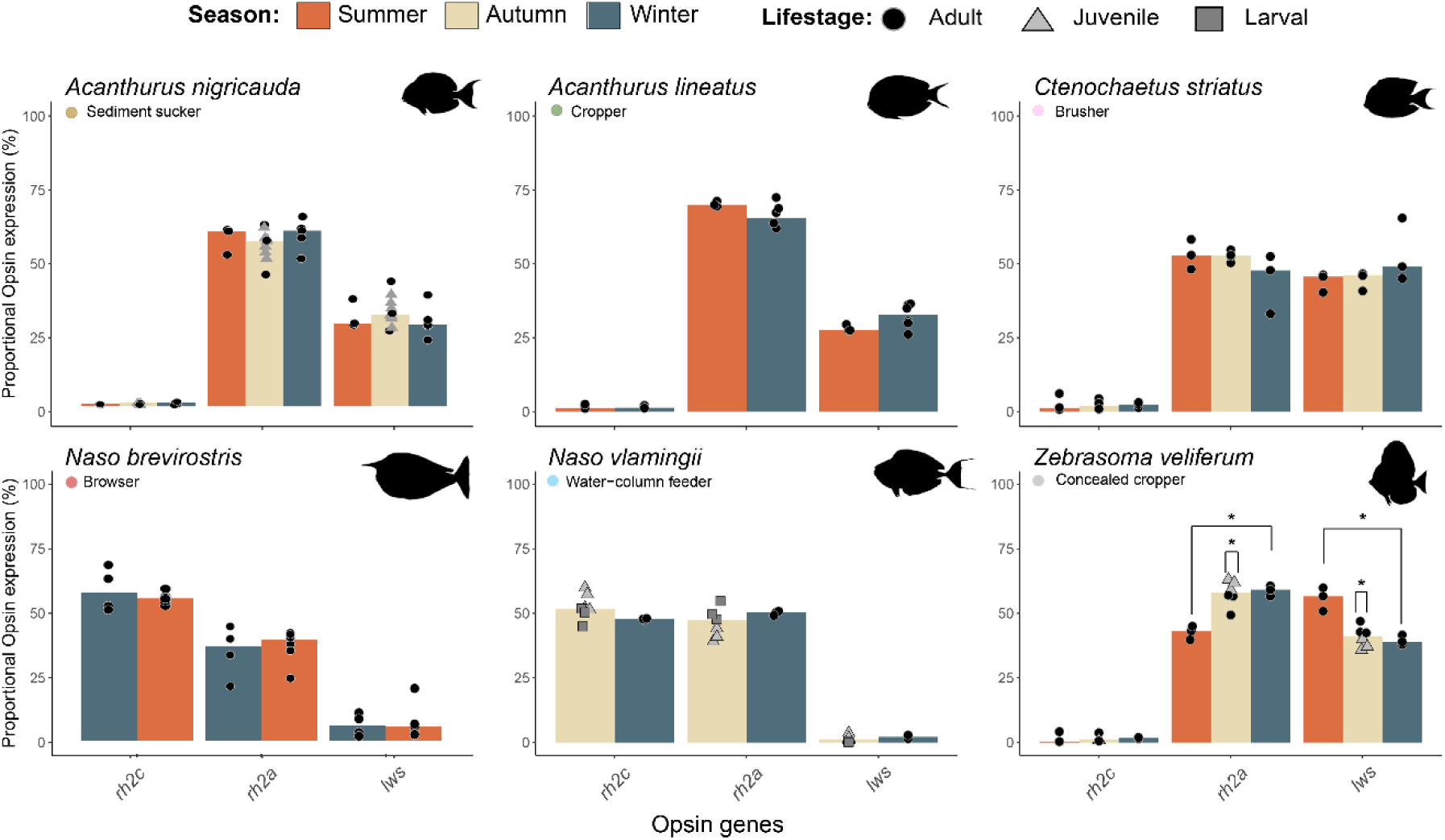
Seasonal and developmental opsin gene expression in surgeonfishes: Proportional double cone opsin (*rh2a, rh2c* and *lws*) gene expression for six species of surgeonfish across seasons (Summer: orange, Autumn: yellow, Winter: blue). The bignose unicornfish (*Naso vlamingii*) expression was analysed in three stages [larval (square), juvenile (triangle) and adult (circle)]. In comparison, only two stages (juvenile, adult) were investigated in the epaulette surgeonfish (*Acanthurus nigricauda)* and the sailfin tang (*Zebrasoma veliferum*). The height of the bars represents the median expression. *Z. veliferum* was the only species showing seasonal differences in *lws* and *rh2a* expression between summer and winter (see supplementary Table S5). Additionally, in autumn, adult and juvenile *Z. veliferum* individuals differed significantly in the expression of their double cone opsin genes. No seasonal difference was found for the single cone expression (*sws2*; see Table S3 for details). Opsin gene expression was compared across seasons and life stages using a Mann-Whitney or Kruskal-Wallis test (*p < 0.05).

## Discussion

Surgeonfishes play a key role on coral reefs, performing functions such as removing macroalgae and feeding on plankton in the water column (Choat et al., 2002; Friedman et al., 2016; Tebbett et al., 2022). Among the 22 surgeonfish species studied, we identified two main opsin expression profiles that are independent of the functional group to which the species belong. Instead, they coincide with the phylogenetic split between the two main subfamilies; Acanthurinae have a longer-shifted visual system compared to the Nasinae (Figs. 1 and 2). Moreover, the phylogenetic inertia in opsin gene expression was strong, with little seasonal or developmental plasticity in our dataset (Figs. 3 and 4). However, while opsin gene expression was mostly fixed, several other retinal genes, including ones related to cell physiology, regulatory processes and cellular morphology, exhibited significant seasonal expression plasticity (Fig. 3).

### Opsin gene expression profiles of surgeonfishes

Consistent with previous investigations into the surgeonfish visual system, the species in this study expressed five cone opsin genes, including *sws2Aα, sws2b, rh2a, rh2c*, and *lws*, with only minor amino acid differences between species (Fogg et al., 2023; Tettamanti et al., 2019), Table S3). Benthic-feeding functional groups belonging to the Acanthurinae subfamily, such as sediment suckers, brushers and croppers, expressed a longer-shifted palette comprising *sws2Aα, rh2a* and *lws*. By comparison, within the Nasine subfamily, such as water-column feeders and, unexpectedly, browsers showed high expression of *sws2b, rh2c*, and *rh2a*, consistent with a shorter-shifted palette (Fig. 2). As suggested by Stieb and colleagues in damselfishes and cichlids (Carleton & Kocher, 2001; Stieb et al., 2023), longer-wavelength sensitivity (i.e, high *lws* expression) is likely to assist benthic feeding and finding far-red-reflecting algae, while shorter-wavelength sensitivity (i.e. *sws1* expression) is helpful when looking for ultraviolet-absorbing or scattering zooplankton and gelatinous prey. For surgeonfishes, although the high expression of the violet-sensitive *sws2b* opsin (estimated λmax ∼ 424 nm, Fig. 1, Table S3 and S4) may extend far enough into the shorter wavelengths to support zooplankton detection (Siebeck & Marshall, 2007), surgeonfishes have a lens transmission cutoff at ∼400 nm (Siebeck & Marshall, 2001). Hence, for surgeonfish, the association between opsin gene expression and functional group or feeding mode is less clear.

Differences between opsin expression and functional group are further seen in blue tang (*P. hepatus*), which displays a long-shifted palette similar to that of benthic-feeding species in the genera *Acanthurus* and *Ctenochaetus* (Fig. 2), despite being a water-column feeder (Friedman et al., 2016). Conversely, within the Nasinae group, the browser species (e.g., *N. unicornis*) that primarily feed on macroalgae (Puk et al., 2016; Topor et al., 2019) share a similar opsin expression profile with their water-column-feeding relatives (e.g., *N. brevirostris*) (Fig. 2). These contrasting opsin expression profiles in surgeonfishes may be largely driven by evolutionary history, as shown by the strong phylogenetic signal (Pagel’s λ = 1) in the relationship between *sws2*, *rh2c*, and *lws* expression and feeding mode. Consequently, present-day opsin expression profiles appear to reflect ancestral functional groups rather than current ecological niches. For instance, the common ancestor of Acanthuridae was a cropper, a foraging strategy previously associated with sensitivity to longer wavelengths (Stieb et al., 2023). In contrast, the ancestor of the Nasinae was a water-column feeder, likely possessing a visual system sensitive to shorter wavelengths (Stieb et al., 2024; Tebbett et al., 2022; Tettamanti et al., 2019). The conserved opsin expression profiles and lack of phenotypic plasticity observed in this study suggest that surgeonfishes visual systems may be more evolutionarily constrained than those of other fish families [e.g., damselfishes (Stieb et al., 2016), and killifish (Fuller et al., 2005)], potentially limiting their capacity to respond to environmental change through either short-term regulatory plasticity or longer-term evolutionary adaptation.

### Developmental opsin gene expression plasticity

In coral reef fishes, significant shifts in opsin gene expression are often correlated with extreme shifts in diet and light habitat, where the larvae migrate from a pelagic, shorter wavelength-dominated blue environment to a broader spectrum, often longer wavelength-dominated greener coral reef environment (Fogg et al., 2022; Marshall et al., 2003; Tettamanti et al., 2019, 2025). Previous studies have shown ontogenetic changes in opsin gene expression in surgeonfishes during this settlement phase, shortly before metamorphosis takes place [convict surgeonfish (*A. triostegus*; Fogg et al., 2023) and spotted unicornfish (*N. brevirostris*: Tettamanti et al., 2019)]. Once settled, our results showed that the shift from juvenile to adult in three surgeonfish species did not differ in opsin expression, similarly to previous research in *N. brevirostris* (Tettamanti et al., 2019) (Fig. 4). Surprisingly, *N. vlamingii,* showed no differences across any of the three life stages that were assessed, from larvae to juveniles to adults. Notably, this species transitions from an open-ocean phase as larvae to deep-lagoon and drop-off habitats as adults, where they feed on zooplankton in midwater aggregations (Randall, 2002). The resulting similarity in diet and light environment (Marshall et al., 2003) may underpin stable opsin gene expression in this case.

### Seasonal flexibility in retinal and opsin gene expression

Our spot measurements of the light environment and data logger measurements from AIMS demonstrate this seasonal shift in the light habitat on the reefs around Lizard Island (Fig. S2). Specifically, the light intensity is much higher in summer compared to winter. The spot measurements also showed a generally broader spectral range in winter, when the water is clearer. Supporting these measurements, long-term data showed substantially higher PAR and a greener spectral environment in summer than in winter, with patterns that overlapped significantly with seasonal temperature changes. The variation observed in daily PAR measurements may be partially explained by the high precipitation in summer, when river runoff brings high loads of sediment and dissolved organic material to inshore reefs along the GBR (Devlin et al., 2001; Schroeder et al., 2012).

Among coral reef fishes, our knowledge of phenotypic flexibility is limited to a few families and encompasses only a small subset of species within each group (Shaughnessy & Cortesi, 2024). Because surgeonfish experience constant fluctuations in light conditions through daily movements within and between reefs, most surgeonfish, as predicted, showed no seasonal change in opsin gene expression (Fig. 3, Table S5). The exception was sailfin tang (*Z. veliferum*), which increased the proportional expression of *lws* in summer and decreased the expression of *rh2a* (Fig. 4). The significance of this shift remains unclear, but *Z. veliferum* may have some feeding plasticity (Letourneur et al., 2017; Tebbett et al., 2022) and thus may switch from a primary algivorous red diet in summer to a detritivorous green/brown diet in winter.

Surprisingly, both *N. unicornis* and *N. brevirostris* showed an upregulation in absolute expression across all their opsin classes in winter. As light intensity and temperature are higher in summer, one might expect opsin turnover and gene expression to increase accordingly. This is somewhat supported by higher overall opsin expression in the summer in many of the other surgeonfish species, such as lined surgeonfish (*A. lineatus*; Table S6). On the other hand, similar to what was found in cichlids (Bolstad & Novales Flamarique, 2023; Wagner & Kröger, 2005), it is possible that the higher opsin expression in *Naso* species is due to an increase in their photoreceptor length to maximise light capture in the dimmer winter months (Fig. S6). Further supporting this hypothesis, *ndrg1-b*, a gene known to be involved in photoreceptor outer segment formation, was upregulated in both *Z. veliferum* and *N. brevirostris* (Fig. 3, Table S8) (Takita et al., 2016). Thus, increasing the outer segment length of cones could be a compensatory mechanism to enhance photon catch and overall sensitivity in low-intensity light environments (Hárosi & Novales Flamarique, 2012).

To date, surgeonfish phenotypic flexibility in colour vision has only been observed in a laboratory setting, and plastic opsin expression in adult *A. triostegus* occurred only in response to extreme differences in the light environment, e.g., constant dim light and constant bright light (Fogg et al., 2023). Given this, the observed seasonal changes in the light environment on the reefs may fall within the natural variation surgeonfish regularly experience. For instance, large non-resident *N. unicornis* adults move between reefs or to deeper, more exposed reef slope habitats in response to high macroalgal densities (Welsh & Bellwood, 2015). This suggests that larger browser surgeonfish taxa can have extensive home ranges with constantly fluctuating light environments (Meyer & Holland, 2005; Welsh & Bellwood, 2015). Therefore, seasonal light environment shifts around Lizard Island may not be significant enough to drive colour-vision flexibility in many surgeonfish species at the molecular level (Fig. 4, Table S5).

While opsin gene expression remained largely unchanged, other retinal genes showed notable changes, many of which were involved in cellular homeostasis rather than colour vision. Notably, genes involved in oxygen carrier activity (GO:0005344) and aerobic respiration (GO:0009060) were upregulated in summer in both *Z. veliferum* and *N. brevirostris* (Fig. 3, Table S7 and S8). Previous studies have shown seasonal increases in oxygen delivery and red blood cell count in fishes, whilst maintaining aerobic scope when the water is warmer (Collazos et al., 1998; Fry & Hart, 1948; Tran & Johansen, 2023). Thus, upregulation of genes involved in aerobic pathways may provide a mechanism to compensate for seasonal temperature effects and maintain visual performance year-round. In addition, the top genes upregulated in the surgeonfishes in summer were involved in pathways such as translation (GO:0006412), RNA binding (GO:0003723), and DNA-binding transcription activity (GO:0000981) (Table S7 and S8). The upregulation of genes involved in these pathways is likely caused by higher temperatures increasing enzymatic activity and DNA flexibility/coiling (Bisswanger, 2017; Driessen et al., 2014).

### Conclusion

Our comparative transcriptomic analyses revealed two major opsin gene expression profiles in surgeonfishes: Nasinae are short-wavelength-shifted compared to Acanthurinae. This divergence, and therefore, by extension, colour vision in surgeonfishes, arose during the ancestral split between the two subfamilies ∼80 Ma and has since been largely conserved throughout their evolutionary history. Moreover, we found that surgeonfish opsin gene expression has very little scope for phenotypic plasticity in development or between seasons. Hence, the dual nature of surgeonfish vision seems ‘good enough’ to exploit a wide range of visual niches. It suggests that, opposed to what is known from smaller territorial species (Stieb et al., 2023, 2024), for larger, highly mobile reef fishes, broad visual tuning may represent an alternative evolutionary strategy for adapting to dynamically changing light environments. With tropical coral reefs at the forefront of rapidly escalating environmental change (Hughes et al., 2003; Vercelloni et al., 2020), understanding how plastic and constrained visual systems influence behaviour will be critical for predicting species resilience and adaptive potential.

## Materials and Methods

### Animal collection and tissue preservation

Adult and juvenile surgeonfishes in this study were caught at depths between 1 and 12 m using various methods, including a diluted clove oil solution (200 ml clove oil per 1 L of seawater), hand nets, barrier nets, and spear guns. Larval individuals were caught using crest nets.

To assess visual specialisation, a phylogenetically representative set of Acanthuridae (22 out of 86 species with representatives from five out of the six genera, n = 108 individuals) was collected from the coral reefs surrounding Lizard Island (northern Great Barrier Reef; 14°40 ′S, 145°27 ′E) and Heron Island (southern Great Barrier Reef; 23°26 ′S, 151°55 ′E), Australia in 2016 – 2018 (Fig. 1, Table S1). To further investigate seasonal flexibility in opsin gene expression, a subset of seven surgeonfish species was selected. For each species, an additional three to five individuals were sampled during both the summer (December – April) and winter (June – August) months between 2020 – 2021 (total n = 43 individuals, Table S1).

In addition, we explored differences in opsin expression across developmental stages in four surgeonfish species: *Acanthurus nigricauda, Naso vlamingii, Zebrasoma scopas,* and *Z. veliferum.* We investigated three life stages for *N. vlamingii*. Specifically, three larval individuals were collected from the reef crest of the lagoon at Temae, Moorea, French Polynesia (17°29’S, 149°45’W) in March 2018, whilst juveniles and adults of the same species were collected from Lizard Island in 2016 and 2018. Two life stages (juvenile and adult) of *A. nigricauda, Z. scopas,* and *Z. veliferum* were also collected from Lizard Island (Table S1).

All animals were anaesthetised using an overdose of clove oil or an ice slurry under approval from the Animal Ethics Committee of The University of Queensland (2022/AE000569 and 2016/AE000304). Dissections were consistently done at 2 pm to align with peak cone opsin gene expression (Yourick et al., 2019). Each individual was photographed, and its standard length (SL) was measured (See Table S1). Individuals were not sexed due to the absence of pronounced reproductive organs. Retinas were carefully dissected from the eyecup and preserved in RNAlater at –80° C for subsequent transcriptome analyses (Table S1). All collections were conducted under the Great Barrier Reef Marine Park Permit (G17/38160.1 and G17/38160.2) and Queensland General Fisheries Permit (207976 and 266344).

### Seasonal light environment measurements

We first quantified differences in the light environments of coral reefs using measurements taken during a summer (December 2021) and winter (July 2021) season. The seasonal light environment measurements were taken at the same position on the lagoon reef at Lizard Island (Fig. S2). The sidewelling absolute irradiance measurements were taken using a USB2000 spectrophotometer running SpectraSuite software (Ocean Optics), in which a cosine-corrected 600μm optic fibre was connected to the spectrometer. The sidewelling irradiance light was taken as we assumed that the fish are mostly viewed from the side by con– and heterospecifics. The optic fibre was positioned 1-2 m underwater above the coral reef whilst connected to a spectrometer on a boat. Five repeated measurements of the light environment were taken and averaged. All measurements were performed mid-day (11-1 pm) on a day with little cloud cover. Each season’s measurements were gathered on a single day. Before the measurements, the spectrophotometer and optic fibre were calibrated using a DH-2000 Deuterium Tungsten Halogen light source (Ocean Optics).

From the Australian Institute of Marine Science (AIMS, https://apps.aims.gov.au/ts-explorer/ Date accessed: 08/2024), we gathered annual light environment and temperature sensor data. These sensors measured the seasonal fluctuations of temperature (°C) and photosynthetically active radiation (PAR; 400-700nm). The average PAR and temperature were calculated for each day from September 2016 to May 2020. Thus, a three-year seasonal cycle could be observed.

### Transcriptome sequencing, quality filtering, and de novo assembly

Total RNA was extracted following the manufacturer’s protocols using a RNeasy Midi Kit (Qiagen) (Acanthuridae). A Eukaryotic Total RNA Nanochip on an Agilent 2100 BioAnalyzer determined the quality and concentration of the extracted RNA. We then outsourced the bulk retinal-transcriptome sequencing (RNAseq) and library preparation [3 Gb/sample (approx. 10 million reads), non-stranded, 150bp paired-end, 300bp insert] to Novogene (https://en.novogene.com/).

The protocol for *de novo* assembly and transcriptome filtering followed the methods described in de Busserolles et al. (2017). In short, the raw reads were uploaded to the Genomics Virtual Laboratory (v.4.0.0) (Afgan et al., 2015) on the Galaxy Australia server (https://usegalaxy.org.au/). The FastQC (Galaxy v.0.72) tool assessed the quality of the sequences, and sequences were then filtered using Trimmomatic (Galaxy v. 0.36.6) (Bolger et al., 2014). Trinity was then used to perform the *de novo* assembly of one reference transcriptome per species (default settings, paired-end, –-min_kmer_cov = 4; Galaxy v.2.4.0.2; Haas et al., 2013).

### Opsin gene mining and phylogenetic reconstruction

To extract the visual opsin gene sequences from the transcriptomes, we followed the protocol outlined by de Busserolles et al. (2017) and Musilova et al. (2019). The assembled transcripts of our focal surgeonfish species were mapped to the opsin genes of previously investigated species, including *Naso brevirostris*, *Acanthurus triostegus* and the dottyback, *Pseudochromis fuscus* (Cortesi et al., 2016; Fogg et al., 2023; Tettamanti et al., 2019). These species were chosen because of overlap or close phylogenetic relationships with the species of interest, with *P. fuscus* serving as a reference for all opsin gene classes. Mapping was done with medium sensitivity (30% max mismatch between transcripts) in Geneious v. 2020.1.2 (www.geneious.com). For raw reads, a medium-low sensitivity was applied against the extracted opsin sequences (20% max mismatch between reads). Mapped reads were visually inspected using single-nucleotide polymorphisms (SNPs) to identify any opsin gene duplicates, which were manually extracted, if necessary, following the method detailed in de Busserolles et al. (2017).

BLASTn (http://blast.ncbi.nlm.nih.gov/) was initially used to confirm the identity of the opsin genes. A phylogenetic analysis of the opsin genes was then performed to assign gene duplicates to their orthologs across different species. The extracted opsin sequences were aligned with a reference data set (obtained from GenBank https://www.ncbi.nlm.nih.gov/genbank, Table S2) using MAFFT v.7.450 (Katoh & Standley, 2013) with the L-INS-I settings. On the CIPRES platform (https://www.phylo.org/), jModeltest v.2.1.6 (Darriba et al., 2012) was used to select the most appropriate model of sequence evolution. Following this, we used MrBayes v.3.2.7a (Ronquist et al., 2012) to infer the phylogenetic relationship between the genes. The following parameters were applied: GTR+I+γ model, with two independent MCMC searches (four chains each), 10 million generations per run, burn-in of 25%, generations, and a tree sampling frequency of 1,000. A majority-rule consensus tree was generated from the sumt files and imported to FigTree 1.4.4 (Rambaut, 2009) for figure creation.

### Proportional opsin gene expression

To quantify the proportional opsin gene expression across seasons and development, the filtered reads were mapped against the coding regions of each species’ opsin repertoire, as per de Busserolles et al. (2017) and Tettamanti et al. (2019). For each opsin gene (*i*), the read fragment count (*R_i_*) was normalised to the length of its coding sequence (*bp_i_*): *R_i_*_,*normalised*_ = (*R_i_*/*bp_i_*).

For the rod opsin, the proportion of *rh1* (*p_i_*_,*rod*_) was divided by the total normalised opsin gene expression (*T_opsin_*).

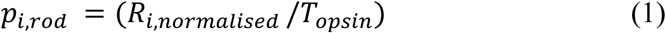

The proportional expression of single cones (*p_i_*_,*SC*_) and double cones (*p_i_*_,*DC*_) was calculated by the proportion of the normalised opsin gene expression (*R_i_*_,*normalised*_) out of the total normalised expression for single (*T_SC_*) or double cones (*T_DC_*). Depending on what type of cone opsin is expressed, i.e., *sws1* and *sws2* in single-, and *rh2* and *lws* in double cones, the two following formulas were used:

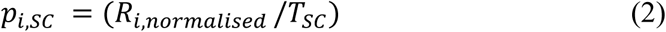

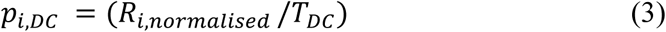

### Relative opsin gene expression

Relative opsin gene expression was determined by normalising opsin expression to two housekeeping (HK) genes. Two stably expressed HK genes, β-actin (*actb*) and elongation factor-1α (*elf1a*), are commonly used to normalise opsin gene expression in fishes (Yourick et al. 2019). The species-specific HK gene sequences were extracted from the transcriptomes by mapping the assembled transcriptomes to previously published coding sequences from the convict surgeonfish, *Acanthurus triostegus* (*actb*, NCBI accession number ON692917; *elf1a*, ON692918).

With a custom sensitivity setting, the transcriptomic reads were mapped against the species-specific HK genes (maximum gap per read, 15%; word length, 14; maximum mismatches per read, 5%; maximum gap size, 50 bp; index word length, 12). The number of mapped reads to HK genes was normalised by the gene length. To calculate the relative opsin gene expression, the normalised reads for each opsin gene were divided by the mean of the normalised reads for the HK genes.

### Statistical analysis

All statistical analyses were conducted using R Studio (version 2023.03.0, R Core Team, 2020). Due to small sample sizes, our data did not meet the assumption of parametric testing and had low statistical power. Thus, to compare the statistical significance of relative and proportional opsin expression between two groups across seasons and development, the data were analysed using one-tailed Mann-Whitney tests. To control for seasonal effects, comparisons of opsin expression across developmental stages were limited to individuals collected within the same season. For comparisons of more than two groups, the data were analysed with a Kruskal-Wallis test and a Dunn’s multiple comparisons test.

### Whole-transcriptome differential gene expression analysis

Differential gene expression analyses were conducted for two species of surgeonfish; a subset of species to investigate which retinal genes, aside from opsin genes, exhibited significant seasonal changes. To quantify transcript abundances in surgeonfishes, a mapping-based approach using Salmon quant was used on the Galaxy platform (version 1.10.1; default settings except specifying paired-end, unstranded reads (U), discarding orphan quasi, validating mappings, and mimicking Bowtie) (Patro et al., 2017). Using Trinity, a reference transcriptome was *de novo* assembled using the reads of one paired-end library from each season. A read count matrix was then uploaded to the iDEP platform (v.2.0 http://bioinformatics.sdstate.edu/idepg) (Ge, 2021), where the samples were pre-processed by removing features of less than 0.5 counts per million and transformed for clustering with a rlog algorithm. A summer *vs*. winter comparison was made using DESeq2 with a p-adjusted value of < 0.05 and a minimum fold-change (FC) of 2. The fold-change was used to assess the top 15 up– and down-regulated genes.

DGE lists were re-formatted to create gene ID-to-sequence lists (FASTA format) in Galaxy. NCBI BLAST+ blastx (Galaxy Version 2.14.1+galaxy2) was used to identify the top BLAST hits against the Swiss-Prot (2023) database, whilst Blast top hit descriptions v.0.1.2 was used to annotate DEGs (Altschul, 1997; Camacho et al., 2009; Cock et al., 2013, 2015). To identify the GO terms and functions of the top 15 up– and downregulated DGEs, UniProtKB and QuickGO were used for manual searches (See Table S9).

### Spectral sensitivity estimations

Spectral sensitivities (λmax) of surgeonfish visual pigments were estimated using the opsin gene sequences identified here. Sequences were translated into amino acid sequences using GENEIOUS and aligned with bovine rhodopsin (NP_001014890.1) using MAFFT (v. 7.222) (Katoh and Kuma 2002). Following the site numbering of amino acids in the bovine rhodopsin sequence allowed the identification of surgeonfish specific amino acid residues that correspond to known tuning and binding pocket sites according to bovine rhodopsin protein structure (Palczewski et al., 2000). Protein sequences were then aligned with reference sequences of well-studied model systems with known visual pigment spectral sensitivities either through in-vitro expression or MSP, and their visual pigment spectral sensitivities were inferred based on sequence differences to each primary reference sequence: RH1: *Oryzias latipes* RH1 (Matsumoto et al. 2006); SWS2B: *Oreochromis niloticus* SWS2B (Parry et al., 2005); SWS2A: *O. niloticus* SWS2A (Parry et al., 2005); RH2B: *O. niloticus* RH2B (Parry et al., 2005) and *Metriaclima zebra* RH2B (Spady et al., 2006); RH2A, *O. niloticus* RH2Aβ (Parry et al., 2005); LWS, *O. niloticus* LWS (Parry et al., 2005). We focused on variable amino acid residues at either known tuning sites or retinal-binding pocket sites. Site effects were then either inferred for known substitutions or for substitutions that cause a change in polarity. Where ambiguous or uncertain site effects did not allow a single estimate, we provided a range of λmax values between an upper and lower limit. To account for high sequence diversity among RH2C opsins, including possibly functional yet untested sites and effects, the upper and lower limit estimates for surgeonfish RH2C visual pigments were calculated using *M. zebra* RH2B (484 nm) and *O. niloticus* RH2B (472 nm), respectively. For a detailed description of the considered site effects, see Supplementary Information.

To further support the amino-acid-based λmax estimations, we also used a regression-based machine learning model integrated into the Visual Physiology Opsin Database (Frazer et al., 2024). We used the ‘whole-dataset’ base model, which incorporates the most extensive dataset available (861 vertebrate opsins), to train the model and further predict the spectral sensitivities of the surgeonfish visual pigments.

### PGLS and PGLM

To examine the evolutionary relationship between double-cone opsin expression and ecological traits (functional group and feeding strategy), we conducted a phylogenetic generalised least squares regression (PGLS) using the R ‘Geiger’ package (Pennell et al., 2014). Since the expression of *sws2Aα* and *sws2b* in single cones within surgeonfish was close to 0% or 100%, a phylogenetic generalised linear model (PGLM) was employed to handle the bimodal distribution with the R ‘phylolm’ package, respectively (R Core Team, 2020; Tung Ho & Ané, 2014). Initially, we categorised species into six functional groups following Tebbett et al. (2022), but then grouped surgeonfish species into three broader trophic categories based on feeding behaviour (herbivores, zooplanktivores, or omnivores) to increase the model’s power. We obtained the most comprehensive surgeonfish phylogeny to date from Sorenson et al., 2013. The PGLS regression estimates the maximum-likelihood (ML) value of the phylogenetic scaling factor λ (Pagel’s λ), where λ = 1 indicates complete phylogenetic dependence and λ = 0 signifies no phylogenetic influence.

## Data accessibility

All data supporting this manuscript, including information for spectral sensitivity estimations, transcriptome assemblies, sequence alignments and R code, are available on UQ eSpace: tba. Transcriptomic raw reads and annotated visual opsin gene sequences are available through GenBank (https://www.ncbi.nlm.nih.gov/genbank/) under BioProject accession number tba (BioSamples: tba) and accession numbers tba, respectively.

## Acknowledgements

We would like to acknowledge the Dingaal, Ngurrumungu, Bailai, Gooreng Gooreng, Gurang, and Taribelang Bunda peoples as the traditional custodians of the land and sea Country of the Lizard Island and Heron Island region, from where specimens were collected. We also acknowledge the Traditional Owners of the land on which the University of Queensland is situated, the Turrbal and Jagera peoples. We pay our respects to their ancestors and their descendants, who continue cultural and spiritual connections to Country and recognise their valuable contributions to Australian and global society. In addition, we would like to thank the staff at Lizard Island Research Station (LIRS) and Heron Island Research Station (HIRS) for their continuous support during fieldwork. We would like to thank Dr Derek Sun, Dr Fanny de Busserolles, Dr Sara Stieb and UQ’s broader Marine Sensory Ecology Group for their assistance with fieldwork, specimen collection, and insightful discussions about the work. We would like to acknowledge the staff of Novogene Co., Ltd. for library preparation and transcriptome sequencing.

## Author contributions

A.S. and F.C. conceived the study. A.S., V.T. and F.C. collected samples for RNA-seq and A.S. and V.T. performed RNA extractions. A.S., V.T., F.C. and M.L. curated and analysed the data. A.S. visualised the data. A.S. wrote the initial manuscript draft, with contributions from M.L. and F.C. All authors commented on and approved the final manuscript.

