## Supplementary Material for "A Deep Phylogenetic Split Between Short- and Long-Wavelength-Tuned Visual Systems in Surgeonfishes (Acanthuridae)"

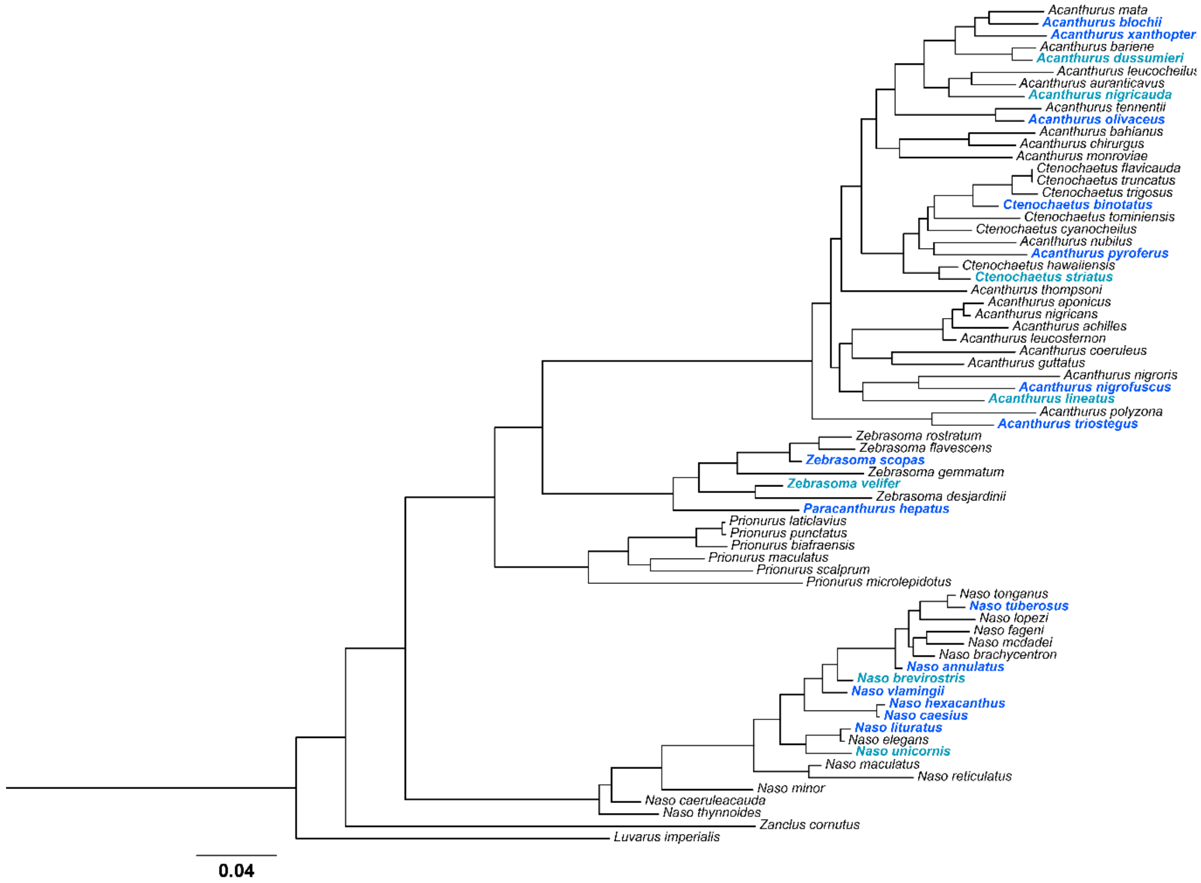


**Figure S1** Phylogenetic tree of the Acanthuridae (surgeonfish) family: Species used in this investigation are highlighted in blue (n = 22), and seven species were used for the seasonal comparative transcriptomics (light blue). The tree was obtained from the Fish Tree of Life (Rabosky et al., 2018). With the exception of *Prionurus* spp. at least one species per major clade and genus was sampled.

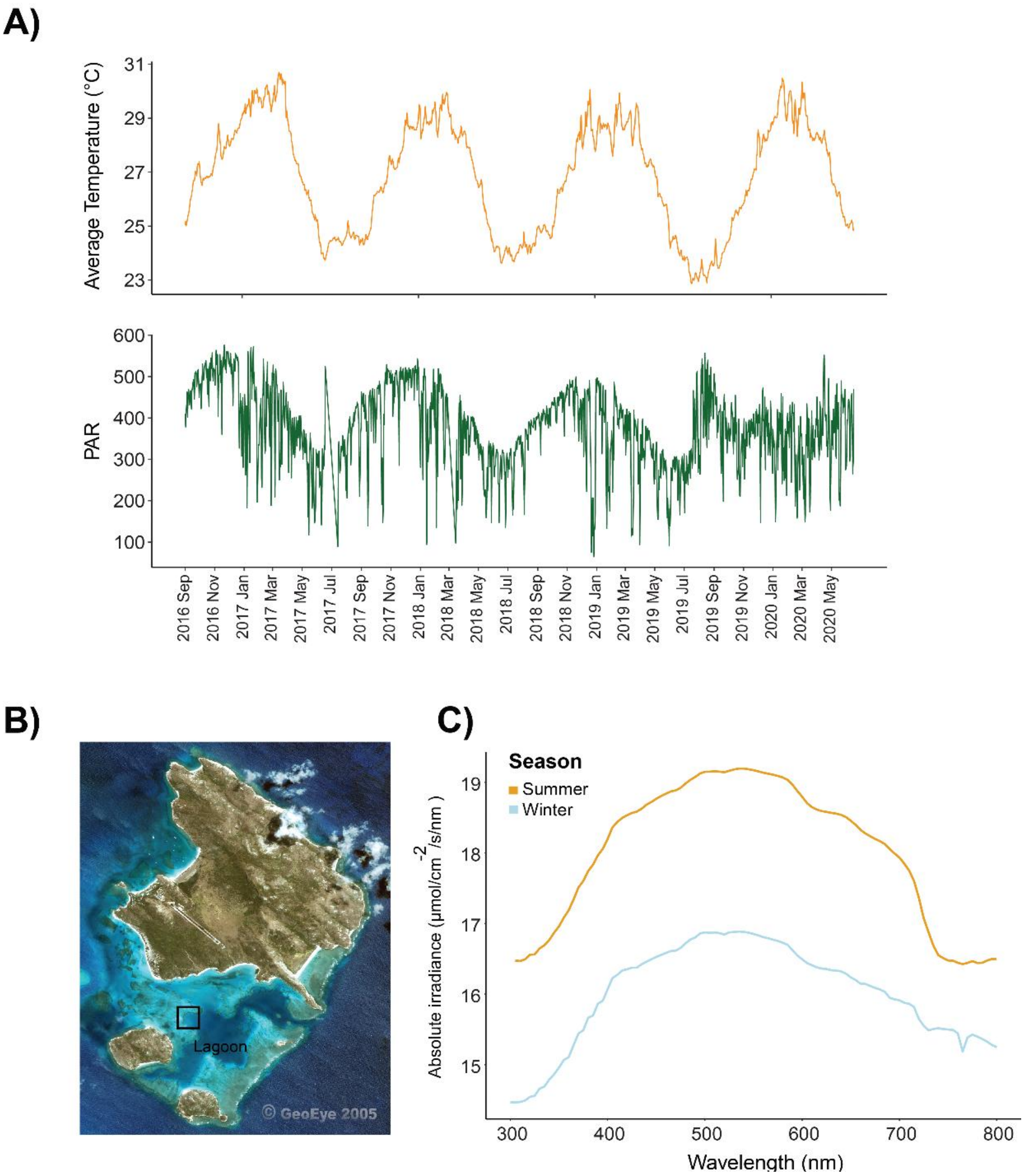


**Figure S2** Seasonal differences in the underwater light environment and temperature at Lizard Island**.** Seasonal fluctuations in (A) average temperature (°C) and PAR (photosynthetic active radiation) from 2016 – 2020 at the main lagoon reefs. The sensors were deployed by the Australian Institute of Marine Science (AIMS), and data was retrieved from AIMS Time Series Explorer (https://apps.aims.gov.au/ts-explorer/ Date accessed: 08/2024). (B) Additional absolute spectral irradiance measurements where taken for each season in the Lizard Island lagoon. (C) Absolute spectral irradiance recorded during summer (yellow) and winter (blue) at a Lizard Island reef. Only side welling irradiance light is shown, representing that the fish are predominately viewed from the side by conspecifics and heterospecifics.

**Supplementary Table 1:** Details of Acanthuridae used in this study. All species were collected from reefs around Lizard Island, Australia, and used for differential gene expression analyses. Surgeonfish were sampled during the summer (December to February), autumn (end of March to May) and winter months (July and August; 3-5 adult individuals per season). Eye size was measured from nasal to temporal (n/t) and dorsal to ventral (d/v) regions of the eye.

| | | | | | | Eye diameter (mm) | |
|---|---|---|---|---|---|---|---|
| **Species** | **Season** | **Lifestage** | **Year** | Location | **Std length (mm)** | **n/t** | **d/v** |
| *Acanthurus blochii* | Autumn | Adult | 2016 | LI | 157.2 | 9.8 | 9.6 |
| | Autumn | Adult | 2016 | LI | 223.0 | 15.0 | 14.0 |
| | Winter | Adult | 2016 | LI | 189.4 | 12.0 | 11.3 |
| | Winter | Adult | 2016 | LI | 130 | 7.0 | 7.4 |
| *Acanthurus dussumieri* | Autumn | Adult | 2016 | LI | 331.2 | 22.5 | 23.5 |
| | Autumn | Adult | 2016 | LI | 170.4 | 13.3 | 12.0 |
| | Autumn | Adult | 2016 | LI | 228.5 | 12.6 | 14.1 |
| | Summer | Adult | 2020 | LI | 273.7 | 14.2 | 15.0 |
| | Summer | Adult | 2020 | LI | 260.1 | 14.6 | 15.1 |
| | Summer | Adult | 2020 | LI | 272.7 | 15.4 | 16.3 |
| | Winter | Adult | 2020 | LI | 287.8 | 14.3 | 15.5 |
| | Winter | Adult | 2020 | LI | 280.0 | 12.8 | 14.7 |
| | Winter | Adult | 2020 | LI | 264.3 | 13.0 | 13.4 |
| | Winter | Adult | 2020 | LI | 267.0 | 13.9 | 15.1 |
| *Acanthurus lineatus* | Summer | Adult | 2020 | LI | 201.4 | 9.6 | 8.7 |
| | Summer | Adult | 2020 | LI | 195.8 | 9.7 | 8.6 |
| | Summer | Adult | 2020 | LI | 201.6 | 11.1 | 8.5 |
| | Winter | Adult | 2016 | LI | 186.1 | 11.9 | 10.8 |
| | Winter | Adult | 2016 | LI | 194.1 | 10.5 | 10.1 |
| | Winter | Adult | 2016 | LI | 161.9 | 7.4 | 8.96 |
| | Winter | Adult | 2021 | LI | 206.5 | 11.4 | 12.4 |
| | Winter | Adult | 2021 | LI | 229.2 | 12.0 | 10.8 |
| | Winter | Adult | 2021 | LI | 206.1 | 9.8 | 9.6 |
| *Acanthurus nigricauda* | Summer | Adult | 2020 | LI | 212.4 | 12.1 | 11.7 |
| | Summer | Adult | 2020 | LI | 200.2 | 10.3 | 10.0 |
| | Summer | Adult | 2021 | LI | 111.2 | 8.4 | 10.6 |
| | Winter | Adult | 2020 | LI | 153.8 | 10.1 | 8.0 |
| | Winter | Adult | 2021 | LI | 142.8 | 8.3 | 8.7 |
| | Winter | Adult | 2021 | LI | 144.5 | 9.3 | 10.5 |
| | Winter | Adult | 2016 | LI | 188.0 | 13.4 | 13.6 |
| | Winter | Adult | 2016 | LI | 235.2 | 13.7 | 12.5 |
| | Autumn | Subadult | 2016 | LI | 126.5 | 8.7 | 9.3 |
| | Autumn | Adult | 2016 | LI | 205.4 | 8.7 | 8.9 |
| | Autumn | Adult | 2016 | LI | 164.4 | 9.3 | 9.0 |
| | Autumn | Juvenile | 2016 | LI | 86.7 | 6.1 | 6.1 |
| | Autumn | Juvenile | 2016 | LI | 77.2 | 6.7 | 6.2 |
| | Autumn | Juvenile | 2016 | LI | 56.9 | 6.1 | 6.3 |
| | Autumn | Juvenile | 2016 | LI | 29.5 | 3.3 | 3.7 |
| | Autumn | Juvenile | 2016 | LI | 27.3 | 3.3 | 2.9 |
| | Autumn | Juvenile | 2016 | LI | 22.8 | 3.2 | 2.9 |

| | | | | | | | |
|---|---|---|---|---|---|---|---|
| | Autumn | Juvenile | 2016 | LI | 22.9 | 2.8 | 3.3 |
| *Acanthurus nigrofuscus* | Autumn | Adult | 2017 | LI | 104.2 | 8.1 | 9.0 |
| | Autumn | Adult | 2017 | LI | 109.8 | 9.1 | 8.7 |
| | Autumn | Adult | 2017 | LI | 107.4 | 7.8 | 8.1 |
| *Acanthurus olivaceus* | Autumn | Adult | 2016 | LI | 232.4 | 10.2 | 11.1 |
| | Autumn | Adult | 2016 | LI | 167.0 | 9.0 | 9.2 |
| *Acanthurus pyroferus* | Autumn | Adult | 2017 | HI | 178.7 | 11.9 | 12.2 |
| | Autumn | Adult | 2017 | HI | 175.3 | 13.0 | 13.4 |
| *Acanthurus triostegus* | Autumn | Adult | 2017 | HI | 61.2 | 5.1 | 5.3 |
| *Acanthurus xanthopterus* | Autumn | Adult | 2017 | HI | 154.4 | 9.6 | 9.8 |
| *Ctenochaetus binotatus* | Winter | Adult | 2016 | LI | 160.1 | 10.9 | 13.1 |
| | Winter | Adult | 2016 | LI | 180.4 | 11.7 | 10.7 |
| | Winter | Adult | 2016 | LI | 73.2 | 7.2 | 6.7 |
| *Ctenochaetus striatus* | Autumn | Adult | 2016 | LI | 147.2 | 10.3 | 10.2 |
| | Autumn | Adult | 2016 | LI | 151.4 | 9.5 | 8.7 |
| | Autumn | Adult | 2016 | LI | 262.4 | 12.1 | 12.5 |
| | Autumn | Subadult | 2016 | LI | 55.7 | 5.4 | 6.3 |
| | Summer | Adult | 2020 | LI | 146.7 | 9.3 | 9.7 |
| | Summer | Adult | 2020 | LI | 175.8 | 9.1 | 9.6 |
| | Summer | Adult | 2020 | LI | 155 | 11.3 | 10.3 |
| | Winter | Adult | 2020 | LI | 146.5 | 9.2 | 7.6 |
| | Winter | Adult | 2020 | LI | 160.9 | 8.5 | 8.1 |
| | Winter | Adult | 2020 | LI | 163.4 | 8.7 | 9.5 |
| *Naso vlamingii* | Autumn | Juvenile | 2017 | HI | 207.3 | 13.7 | 14.6 |
| | Autumn | Juvenile | 2016 | LI | 155.3 | 12.6 | 12.1 |
| | Autumn | Juvenile | 2018 | LI | 189.6 | 7.9 | 8.6 |
| | Autumn | Juvenile | 2018 | LI | 168.9 | 11.1 | 11.7 |
| | Autumn | Larval | 2018 | FP | 31.8 | 3.6 | 3.9 |
| | Autumn | Larval | 2018 | FP | 31.9 | 3.9 | 4.2 |
| | Autumn | Larval | 2018 | FP | 31.1 | 4.1 | 4.0 |
| | Winter | Adult | 2016 | LI | 370.4 | 16.6 | 17.5 |
| | Winter | Adult | 2016 | LI | 317.4 | 18.3 | 21.7 |
| | Winter | Adult | 2016 | LI | 318.8 | 16.9 | 18.7 |
| *Naso annulatus* | Autumn | Adult | 2016 | LI | 198.7 | 15.4 | 14.5 |
| | Autumn | Juvenile | 2016 | LI | 31.02 | 4.7 | 4.5 |
| | Autumn | Juvenile | 2016 | LI | 585.8 | 5.2 | 6.35 |
| *Naso brevirostris* | Summer | Adult | 2018 | LI | 321.6 | 16.2 | 14.8 |
| | Summer | Adult | 2018 | LI | 310.8 | 16.7 | 13.5 |
| | Summer | Adult | 2018 | LI | 242.9 | 15.4 | 12.4 |
| | Summer | Adult | 2020 | LI | 203.2 | 13.8 | 11.5 |
| | Summer | Adult | 2020 | LI | 162.6 | 13.9 | 12.2 |
| | Summer | Adult | 2020 | LI | 146.2 | 12.5 | 12.0 |
| | Winter | Adult | 2020 | LI | 160.2 | 11.8 | 11.6 |
| | Winter | Adult | 2020 | LI | 141.7 | 10.7 | 11.1 |
| | Winter | Adult | 2020 | LI | 157.9 | 10.7 | 10.7 |
| | Winter | Adult | 2020 | LI | 197.7 | 12.8 | 12.4 |
| *Naso caesius* | Winter | Adult | 2017 | HI | 280.9 | 16.6 | 14.5 |
| *Naso hexacanthus* | Winter | Adult | 2017 | HI | 268.6 | 14.1 | 13.5 |
| | Winter | Adult | 2017 | HI | 527.7 | 26.0 | 23.3 |
| *Naso lituratus* | Winter | Adult | 2016 | LI | 187.4 | 12.6 | 11.6 |
| | Winter | Adult | 2016 | LI | 218.1 | 13.5 | 13.3 |
| | Winter | Adult | 2017 | HI | 149.7 | 9.5 | 9.3 |

| | | | | | | | |
|---|---|---|---|---|---|---|---|
| *Naso tuberosus* | Autumn | Adult | 2018 | LI | 417.4 | 21.8 | 21.6 |
| | Winter | Adult | 2017 | HI | 214.7 | 16.9 | 17.8 |
| *Naso unicornis* | Autumn | Adult | 2016 | LI | 129.2 | 11.5 | 11.5 |
| | Autumn | Adult | 2018 | LI | 156.8 | 13.0 | 12.5 |
| | Summer | Adult | 2020 | LI | 156.4 | 11.5 | 10.8 |
| | Summer | Adult | 2020 | LI | 174.9 | 11.9 | 10.5 |
| | Summer | Adult | 2020 | LI | 133.1 | 10.5 | 9.4 |
| | Winter | Adult | 2016 | LI | 108.5 | 10.5 | 10.6 |
| | Winter | Adult | 2021 | LI | 144.9 | 11.7 | 12.9 |
| | Winter | Adult | 2021 | LI | 129.2 | 10.6 | 7.7 |
| | Winter | Adult | 2021 | LI | 134.7 | 12.6 | 13.2 |
| *Paracanthurus hepatus* | Winter | Adult | 2017 | CM | 179.5 | 7.9 | 8.7 |
| | Winter | Adult | 2017 | CM | 161.5 | 6.3 | 7.3 |
| | Winter | Adult | 2017 | CM | 144.9 | 6.3 | 7.9 |
| *Zebrasoma scopas* | Autumn | Adult | 2016 | LI | 79.5 | 8.2 | 7.8 |
| | Autumn | Adult | 2016 | LI | 70.1 | 7.0 | 7.1 |
| | Autumn | Adult | 2016 | LI | 110.0 | 10.0 | 9.5 |
| | Autumn | Juvenile | 2016 | LI | 36.8 | 4.9 | 4.7 |
| | Autumn | Juvenile | 2016 | LI | 27.6 | 4.7 | 4.5 |
| *Zebrasoma veliferum* | Autumn | Adult | 2016 | LI | 131.3 | 9.1 | 10.3 |
| | Autumn | Adult | 2016 | LI | 90.3 | 7.1 | 7.2 |
| | Autumn | Adult | 2016 | LI | 113.9 | 8.9 | 9.3 |
| | Autumn | Juvenile | 2016 | LI | 46.9 | 6.4 | 6.2 |
| | Autumn | Juvenile | 2016 | LI | 44.19 | 6.7 | 5.5 |
| | Autumn | Juvenile | 2016 | LI | 42.8 | 4.9 | 5.52 |
| | Summer | Adult | 2020 | LI | 62.9 | 7.3 | 7.2 |
| | Summer | Adult | 2020 | LI | 197.4 | 11.6 | 11.9 |
| | Summer | Adult | 2020 | LI | 188.4 | 11.3 | 10.9 |
| | Winter | Adult | 2020 | LI | 122.8 | 10.0 | 9.5 |
| | Winter | Adult | 2020 | LI | 170.8 | 9.4 | 9.7 |
| | Winter | Adult | 2020 | LI | 117.9 | 8.4 | 9.3 |

**Supplementary Table 2:** List of species used as the reference dataset for the phylogenetic analysis of the surgeonfish opsin genes. The full opsin gene sequences are available from GenBank. VA opsins are vertebrate ancient opsins. *rh1*= rhodopsin, *sws1* = short wavelength–sensitive type 1, *sws2* = short wavelength–sensitive type 2, *rh2* = middle wavelength-sensitive class 2, and M/*lws* = middle to long wavelength–sensitive.

| Species | Opsin gene | Accession number |
|---|---|---|
| *Anolis carolinensis* | *lws* | XM008103916 |
| *Anolis carolinensis* | *rh1* | NM001291387 |
| *Anolis carolinensis* | *rh2* | AH007735 |
| *Anolis carolinensis* | *rh2ac* | AH004781 |
| *Anolis carolinensis* | *sws1* | AH007736 |
| *Anolis carolinensis* | *sws2* | AF133907 |
| *Anolis carolinensis* | VAopsin | XM008106312 |
| *Callorhinchus milii* | *lws1* | EF565165 |
| *Callorhinchus milii* | *lws2* | EF565166 |
| *Callorhinchus milii* | *rh1* | EF565167 |
| *Callorhinchus milii* | *rh2* | EF565168 |
| *Caranx malampygus* | *sws2Aa* | SRX360285 |
| *Caranx malampygus* | *sws2ab* | SRX360285 |
| *Caranx malampygus* | *sws2b* | SRX360285 |
| *Cyprinus carpio* | *lws* | AB055656 |
| *Cyprinus carpio* | *rh1* | Z71999 |
| *Cyprinus carpio* | *rh2_1* | AB110602 |
| *Cyprinus carpio* | *rh2_2* | AB110603 |
| *Cyprinus carpio* | *sws1* | AB113669 |
| *Cyprinus carpio* | *sws2* | AB113668 |
| *Cyprinus carpio* | VAopsin | AF233520 |
| *Danio rerio* | *lws1* | NM001313715 |
| *Danio rerio* | *lws2* | NM001002443 |
| *Danio rerio* | *rh1_1* | NM131084 |
| *Danio rerio* | *rh1_2* | HM367062 |
| *Danio rerio* | *rh2_1* | AF109369 |
| *Danio rerio* | *rh2_2* | NM182891 |
| *Danio rerio* | *rh2_3* | NM182892 |
| *Danio rerio* | *rh2_4* | AF109370 |
| *Danio rerio* | *sws1* | AF109373 |
| *Danio rerio* | *sws2* | AF109372 |
| *Danio rerio* | VAopsin1 | NM131586 |
| *Danio rerio* | VAopsin2 | NM001110280 |
| *Echeneis naucrates* | *rh2* | XM_029503145.1 |
| *Gallus gallus* | *lws* | NM205440 |
| *Gallus gallus* | *rh2* | M92038 |
| *Gallus gallus* | *sws1* | M92039 |
| *Gallus gallus* | *sws2* | NM205517 |

| | | |
|---|---|---|
| *Gasterosteus aculeatus* | *lws* | KC594699 |
| *Gasterosteus aculeatus* | *rh1* | KC774627 |
| *Gasterosteus aculeatus* | *rh2* | KC594702 |
| *Gasterosteus aculeatus* | *sws1* | KC774626 |
| *Gasterosteus aculeatus* | *sws2Ab* | KC774625 |
| *Geotria australis* | *lws* | AY366491 |
| *Geotria australis* | *rha* | AY366493 |
| *Geotria australis* | *rhb* | AY366494 |
| *Geotria australis* | *sws1* | AY366495 |
| *Geotria australis* | *sws2* | AY366492 |
| *Homo sapiens* | *lws* | NM020061 |
| *Homo sapiens* | *MWS* | NM000513 |
| *Homo sapiens* | *rh1* | NM000539 |
| *Homo sapiens* | *sws1* | NM001708 |
| *Naso brevirostris* | *sws2b* | MN720499 |
| *Naso brevirostris* | *lws* | MN720503 |
| *Naso brevirostris* | *rh2a* | MN720501 |
| *Naso brevirostirs* | *rh2b* | MN720500 |
| *Naso_brevirostris* | *rh1* | MN720502 |
| *Oreochromis niloticus* | *lws* | AF247128 |
| *Oreochromis niloticus* | *rh1* | AY775108 |
| *Oreochromis niloticus* | *rh2aa* | JF262086 |
| *Oreochromis niloticus* | *rh2ab* | JF262086 |
| *Oreochromis niloticus* | *rh2B* | JF262086 |
| *Oreochromis niloticus* | *sws1* | JF262087 |
| *Oreochromis niloticus* | *sws2a* | AF247116 |
| *Oreochromis niloticus* | *sws2b* | AF247120 |
| *Oryzias latipes* | *lwsa* | AB223051 |
| *Oryzias latipes* | *lwsb* | AB223052 |
| *Oryzias latipes* | *rh1* | AB180742 |
| *Oryzias latipes* | *rh2a* | AB223053 |
| *Oryzias latipes* | *rh2b* | AB223054 |
| *Oryzias latipes* | *rh2c* | AB223055 |
| *Oryzias latipes* | *sws1* | AB223058 |
| *Oryzias latipes* | *sws2a* | AB223056 |
| *Oryzias latipes* | *sws2b* | AB223057 |
| *Oryzias latipes* | VAopsin | NM001136515 |
| *Pseudochromis fuscus* | *lws* | ERX1545070 |
| *Pseudochromis fuscus* | *rh1* | ERX1545070 |
| *Pseudochromis fuscus* | *rh2aalpha* | ERX1545070 |
| *Pseudochromis fuscus* | *rh2abeta* | ERX1545070 |
| *Pseudochromis fuscus* | *rh2B* | ERX1545070 |
| *Pseudochromis fuscus* | *sws1* | ERX1545070 |
| *Pseudochromis fuscus* | *sws2Aalpha* | ERX1545070 |

| | | |
|---|---|---|
| *Pseudochromis fuscus* | *sws2Abeta* | ERX1545070 |
| *Pseudochromis fuscus* | *sws2B* | ERX1545070 |
| *Rhinecanthus aculeatus* | *rh1* | ON060693 |
| *Rhinecanthus aculeatus* | *rh2a* | ON060694.1 |
| *Rhinecanthus aculeatus* | *sws2B* | ON060701 |
| *Rhinecanthus aculeatus* | *lws* | ON060692.1 |
| *Rhinecanthus aculeatus* | *rh2B* | ON060695.1 |
| *Rhinecanthus aculeatus* | *rh2c-1* | ON060696.1 |
| *Rhinecanthus aculeatus* | *rh2c-2* | ON060697.1 |
| *Rhinecanthus aculeatus* | *rh2c-3* | ON060698.1 |
| *Rhinecanthus aculeatus* | *rh2c-4* | ON060699.1 |
| *Rhinecanthus aculeatus* | *rh2c-5* | ON060700.1 |
| *Salmo salar* | *lws* | NM001123705 |
| *Salmo salar* | *rh1* | NM001123537 |
| *Salmo salar* | *rh1like* | NM001123536 |
| *Salmo salar* | *rh2* | AY214132 |
| *Salmo salar* | *sws1* | AY214133 |
| *Salmo salar* | *sws2* | NM001123706 |
| *Salmo salar* | VAopsin | SSAF001499 |
| *Seriola dumerili* | *rh2_01* | XM_022746898.1 |
| *Seriola_dumerili* | *rh2-02* | XM_022746739.1 |
| *Seriola dumerili* | *rh2-03* | XM_022746738.1 |
| *Seriola dumerili* | *rh2-04* | XM_022746838.1 |
| *Sparus aurata* | *rh2-03* | XM_030421531.1 |
| *Sparus aurata_* | *rh2-04* | XM_030421138.1 |
| *Sphaeramia orbicularis* | *rh2_01* | XM_030133850.1 |
| *Sphaeramia orbicularis* | *rh2-02* | XM_030133847.1 |
| *Takifugu rubripes* | *lws* | AY598942 |
| *Takifugu rubripes* | *sws2* | AY598947 |
| *Takifugu rubripes* | *rh1* | NM001078631 |
| *Takifugu rubripes* | *rh2* | AF226989 |
| *Xenopus laevis* | *lws* | NM001090645 |
| *Xenopus laevis* | *rh1* | NM001087048 |
| *Xenopus laevis* | *sws1* | NM001085652 |
| *Xenopus laevis* | *sws2* | BC080123 |

**Supplementary Table 3:** Comparative relative opsin gene expression in 21 adult surgeonfish species. Expression levels (%) are shown as a proportion within single (*sws1* and *sws2b*) and double cones (*rh2c, rh2a, lws*). Rod-opsin expression is expressed as a proportion of the total opsin expression. Mean ± s.d.

| | | Single cones | | Double cones | | | Rods |
|---|---|---|---|---|---|---|---|
| Species | Sample size (n) | *sws2a* | *sws2b* | *rh2c* | *rh2a* | *lws* | *rh1* |
| *Acanthurus xanthopterus* | 1 | 100.0 | 0.0 | 2.7 | 58.9 | 38.5 | 87.32 |
| *Acanthurus blochii* | 4 | 100.0 | 0.0 | 2.0 ± 0.9 | 56.9 ± 2.5 | 41.1 ± 2.6 | 83.9 ± 9.03 |
| *Acanthurus dussumieri* | 10 | 100.0 | 0.0 | 3.6 ± 2.2 | 67.0 ± 7.6 | 29.4 ± 7.1 | 85.5 ± 12.6 |
| *Acanthurus lineatus* | 9 | 100.0 | 0.0 | 1.5 ± 0.5 | 67.5 ± 3.9 | 31.0 ± 3.9 | 78.5 ± 3.4 |
| *Acanthurus nigricauda* | 11 | 99.9 ± 0.1 | 0.08 ± 0.1 | 1.3 ± 0.5 | 64.2 ± 6.6 | 34.5 ± 6.7 | 80.1 ± 5.8 |
| *Acanthurus nigrofuscus* | 3 | 100.0 | 0.0 | 0.9 ± 0.1 | 57.8 ± 12.4 | 41.4 ± 12.5 | 73.7 ± 2.0 |
| *Acanthurus olivaceus* | 2 | 100.0 | 0.0 | 2.3 ± 0.4 | 64.9 ± 6.8 | 32.8 ± 6.4 | 83.0 ± 1.7 |
| *Acanthurus pyroferus* | 2 | 100.0 | 0.0 | 4.4 ± 3.6 | 65.1 ± 3.7 | 30.4 ± 0.1 | 89.9 ± 7.5 |
| *Acanthurus triostegus* | 1 | 99.9 | 0.1 | 1.6 | 52.8 | 45.6 | 86.3 |
| *Ctenochaetus binotatus* | 3 | 100.0 | 0.0 | 4.2 ± 1.1 | 67.0 ± 2.2 | 28.8 ± 1.8 | 88.2 ± 3.05 |
| *Ctenochaetus striatus* | 10 | 100.0 | 0.0 | 2.4 ± 1.8 | 50.4 ± 6.8 | 47.2 ± 7.0 | 81.9 ± 5.5 |
| *Naso annulatus* | 3 | 0.0 | 100.0 | 49.3 ± 4.7 | 50.5 ± 4.6 | 0.2 ± 0.1 | 76.4 ± 12.8 |
| *Naso brevirostris* | 10 | 0 | 100.0 | 57.0 ± 5.4 | 36.3 ± 8.3 | 6.7 ± 5.6 | 82.9 ± 6.9 |
| *Naso caesius* | 1 | 0.0 | 100.0 | 56.6 | 32.2 | 11.2 | 92.89 |

| | | | | | | | |
|---|---|---|---|---|---|---|---|
| *Naso hexcanthus* | 2 | 0.0 | 100.0 | 59.0 ± 4.1 | 34.7 ± 4.4 | 6.3 ± 0.3 | 96.7 ± 1.03 |
| *Naso lituratus* | 3 | 0.0 | 100.0 | 64.4 ± 15.6 | 35.4 ± 15.5 | 0.2 ± 0.1 | 90.9 ± 8.9 |
| *Naso tuberosus* | 2 | 0.0 | 100.0 | 47.7 ± 1.6 | 52.0 ± 1.7 | 0.3 ± 0.1 | 96.6 ± 3.4 |
| *Naso unicornis* | 10 | 0.0 | 100.0 | 56.9 ± 5.6 | 38.0 ± 9.3 | 5.1 ± 6.5 | 89.7 ± 4.4 |
| *Naso vlamingii* | 3 | 0.0 | 0.0 | 47.7 ± 0.3 | 50.1 ± 0.9 | 2.1 ± 0.7 | 86.1 ± 7.5 |
| *Paracanthurus hepatus* | 3 | 100.0 | 0.0 | 0.3 ± 0.2 | 64.3 ± 5.3 | 35.4 ± 5.1 | 68.5 ± 1.5 |
| *Zebrasoma scopas* | 3 | 100.0 | 0.0 | 0.9 ± 0.4 | 54.9 ± 10.0 | 44.2 ± 10.3 | 91.7 ± 1.1 |
| *Zebrasoma veliferum* | 9 | 100.0 | 0.0 | 1.7 ± 1.1 | 56.5 ± 3.8 | 41.2 ± 3.2 | 81.2 ± 7.4 |

**Supplementary Table 4:** Surgeonfish visual pigment spectral sensitivities estimated from cone opsin (SWS2B, SWS2A, RH2C, RH2A, LWS) and rod opsin (RH1) amino acid sequences, and measured by microspectrophotometry (MSP) as retrieved from literature (Losey et al. 2003).

**RH1**

| Species | VPOD | AA-modelling | MSP |
|---|---|---|---|
| *A. blochii* | 493.2 | 496 | |
| *A. dussumieri* | 493.2 | 496 | |
| *A. lineatus* | 493.2 | 496 | |
| *A. nigrocauda* | 493.2 | 496 | |
| *A. nigrofuscus* | 493.2 | 496 | |
| *A. olivaceus* | 493.4 | 496 | |
| *A. pyroferus* | 493.4 | 494 | |
| *A. triostegus* | 494.5 | 496 | 493 |
| *A. xanthopterus* | 493.2 | 496 | |
| *C. binotatus* | 482.7 | 484 | |
| *C. striatus* | 492.4 | 494 | |
| *N. annulatus* | 495.2 | 496 | |
| *N. brevirostris* | 493.4 | 496 | |
| *N. caesius* | 493.4 | 496 | |
| *N. hexacanthus* | 493.3 | 494 | |
| *N. lituratus* | 493.4 | 496 | 497 |
| *N. tuberosus* | 495.2 | 496 | |
| *N. unicornis* | 493.4 | 496 | 494 |
| *N. vlamingii* | 493.4 | 496 | |
| *P. hepatus* | 493.5 | 496 | |
| *Z. scopas* | 492.4 | 494 | |
| *Z. veliferum* | 493.2 | 496 | 494 |

**SWS2B**

| Species | VPOD | AA-modelling | MSP |
|---|---|---|---|
| *A. nigricauda* | 426.2 | 424 | |
| *A. triostegus* | 424.7 | 424 | |
| *N. annulatus* | 414 | 424 | |
| *N. brevirostris* | 414 | 424 | |
| *N. caesius* | 414 | 424 | |
| *N. hexacanthus* | 414.2 | 424 | |
| *N. lituratus* | 414.4 | 424 | 421 |
| *N. tuberosus* | 414.2 | 424 | |
| *N. unicornis* | 414 | 424 | 416 |
| *N. vlamingii* | 414 | 424 | |

**SWS2A**

| Species | VPOD | AA-modelling | MSP |
|---|---|---|---|
| *A. blochii* | 444.8 | 452-460 | |
| *A. dussumieri* | 445.9 | 452-460 | |
| *A. lineatus* | 445.9 | 452-460 | |
| *A. nigrocauda* | 445.9 | 452-460 | |
| *A. nigrofuscus* | 445.9 | 452-460 | |
| *A. olivaceus* | 437.4 | 444-454 | |
| *A. pyroferus* | 436.7 | 444-454 | |
| *A. triostegus* | 444.5 | 452-460 | 458 |
| *A. xanthopterus* | 438.5 | 446-454 | |
| *C. binotatus* | 436.8 | 444-454 | |
| *C. striatus* | 444.8 | 452-460 | |
| *P. hepatus* | 421.9 | 444-448 | |
| *Z. scopas* | 445.9 | 456-464 | |
| *Z. veliferum* | 444.8 | 456-464 | 448 |

**RH2C**

| Species | VPOD | AA-modelling | MSP |
|---|---|---|---|
| *A. blochii* | 490.9 | 475-484 | |
| *A. dussumieri* | 490.9 | 475-484 | |
| *A. lineatus* | 491 | 473-482 | |
| *A. nigrocauda* | 491 | 475-484 | |
| *A. nigrofuscus* | 497.9 | 471-480 | |
| *A. olivaceus* | 489.7 | 472-481 | |
| *A. pyroferus* | 489.7 | 475-484 | |
| *A. triostegus* | 489.7 | 470-479 | |
| *A. xanthopterus* | 490.9 | 475-484 | |
| *C. binotatus* | 487.8 | 473-482 | |
| *C. striatus* | 492.3 | 472-481 | |
| *N. annulatus* | 492.9 | 467-476 | |
| *N. brevirostris* | 492.4 | 471-480 | |
| *N. caesius* | 492.9 | 472-481 | |
| *N. hexacanthus* | 492.1 | 471-480 | |
| *N. lituratus* | 492.9 | 470-479 | 496, 521 |
| *N. tuberosus* | 492.9 | 467-476 | |
| *N. unicornis* | 492.9 | 472-481 | 486, 507 |
| *N. vlamingii* | 492.4 | 471-480 | |
| *P. hepatus* | 491 | 472-481 | |
| *Z. scopas* | 491 | 473-482 | |
| *Z. veliferum* | 491 | 473-482 | 511, 523, 517 |

**RH2A**

| Species | VPOD | AA-modelling | MSP |
|---|---|---|---|
| *A. blochii* | 516.5 | 511-513 | |
| *A. dussumieri* | 516.5 | 511-513 | |
| *A. lineatus* | 516.5 | 511-513 | |
| *A. nigrocauda* | 516.5 | 511-513 | |
| *A. nigrofuscus* | 516.5 | 511-513 | |
| *A. olivaceus* | 516.5 | 511-513 | |
| *A. pyroferus* | 516.6 | 511-513 | |
| *A. triostegus* | 516.5 | 511-513 | 518 |
| *A. xanthopterus* | 516.5 | 511-513 | |
| *C. binotatus* | 515.2 | 510-513 | |
| *C. striatus* | 516.5 | 511-513 | |
| *N. annulatus* | 517.2 | 514-516 | |
| *N. brevirostris* | 518.6 | 514-516 | |
| *N. caesius* | 518.6 | 514-516 | |
| *N. hexacanthus* | 517.2 | 514-516 | |
| *N. lituratus* | 518.6 | 514-516 | 495, 521 |
| *N. tuberosus* | 517.2 | 514-516 | |
| *N. unicornis* | 518.6 | 514-516 | 486, 507 |
| *N. vlamingii* | 517.2 | 514-516 | |
| *P. hepatus* | 511.6 | 513-516 | |
| *Z. scopas* | 516.5 | 511-513 | |
| *Z. veliferum* | 516.5 | 511-513 | 511, 523, 517 |

**LWS**

| Species | VPOD | AA-modelling | MSP |
|---|---|---|---|
| *A. blochii* | 546.7 | 554-560 | |
| *A. dussumieri* | 546.7 | 554-560 | |
| *A. lineatus* | 550.2 | 554-560 | |
| *A. nigrocauda* | 546.7 | 554-560 | |
| *A. nigrofuscus* | 550.2 | 554-560 | |
| *A. olivaceus* | 546.7 | 554-560 | |
| *A. pyroferus* | 542.1 | 547-553 | |
| *A. triostegus* | 550.2 | 554-560 | 518 |
| *A. xanthopterus* | 546.7 | 554-560 | |
| *C. binotatus* | 546.7 | 554-560 | |
| *C. striatus* | 546.7 | 554-560 | |
| *N. annulatus* | 547.4 | 554-560 | |
| *N. brevirostris* | 547.4 | 554-560 | |
| *N. caesius* | 546.7 | 554-560 | |
| *N. hexacanthus* | 547.4 | 554-560 | |
| *N. lituratus* | 517.3 | 538-544 | 495, 521 |
| *N. tuberosus* | 547.4 | 554-560 | |
| *N. unicornis* | 530.9 | 538-544 | 486, 507 |
| *N. vlamingii* | 530.9 | 538-544 | |
| *P. hepatus* | 546.7 | 554-560 | |
| *Z. scopas* | 546.7 | 554-560 | |
| *Z. veliferum* | 546.7 | 554-560 | 511, 517, 523 |

**Supplementary Table 5:** Comparative proportional opsin gene expression between winter (July-August), autumn (April - May) and summer (December – March) in seven adult surgeonfish species. Expression levels (%) are shown as a proportion within single (*sws1* and *sws2b*) and double cones (*rh2b, rh2a, lws*). Rod-opsin expression is expressed as a proportion of the total opsin expression. Mean ± s.d.

| | | | Single cones | | Double cones | | | Rods |
|---|---|---|---|---|---|---|---|---|
| Species | Sample size (n) | Season | *sws2a* | *sws2b* | *rh2c* | *rh2a* | *lws* | *rh1* |
| *Acanthurus nigricauda* | 3 | Summer | 99.9 ± 0.05 | 0.04 ± 0.2 | 0.9 ± 0.2 | 64.3 ± 3.4 | 34.7 ± 5.6 | 79.3 ± 4.8 |
| | 3 | Autumn | 99.9 ± 0.2 | 0.2 ± 0.2 | 1.3 ± 0.4 | 61.3 ± 8.1 | 37.4 ± 9.6 | 86.4 ± 5.5 |
| | 5 | Winter | 99.9 ± 0.04 | 0.05 ± 0.04 | 1.5 ± 0.6 | 65.9 ± 6.0 | 32.6 ± 6.2 | 76.8 ± 3.7 |
| *Ctenochaetus striatus* | 3 | Summer | 100 | | 2.8 ±1.7 | 53.1 ± 2.9 | 44.1 ± 1.9 | 82.7 ± 4.1 |
| | 4 | Autumn | 100 | | 2.3 ± 1.7 | 52.7 ± 1.8 | 45.0 ± 2.8 | 85.1 ± 4.2 |
| | 3 | Winter | 100 | | 2.3 ± 0.5 | 44.4 ± 5.8 | 53.1 ± 6.3 | 76.9 ± 0.6 |
| *Naso brevirostris* | 6 | Summer | | 100 | 55.6 ± 2.2 | 37.2 ± 7.2 | 7.2 ± 6.7 | 86.0 ± 6.2 |
| | 4 | Winter | | 100 | 61.5 ±4.7 | 33.2 ± 7.2 | 5.3 ± 2.8 | 78.8 ± 1.2 |
| *Zebrasoma veliferum* | 3 | Summer | | 100 | 1.6 ± 1.3 | 42.6 ± 1.5 | 55.8 ± 2.7 | 81.6 ± 4.8 |
| | 3 | Autumn | | 100 | 1.6 ± 1.8 | 54.3 ± 4.3 | 44.0 ± 2.5 | 80.7 ± 9.2 |
| | 3 | Winter | | 100 | 1.7 ± 0.1 | 58.7 ± 1.2 | 39.5 ± 1.1 | 81.2 ± 4.6 |
| *Naso unicornis* | 3 | Summer | | 100 | 60.0 ± 2.0 | 39.5 ± 1.8 | 0.4 ± 0.3 | 92.3 ± 2.5 |
| | 2 | Autumn | | 100 | 55.1 ± 5.7 | 44.7 ± 5.6 | 0.3 ± 0.2 | 90.2 ± 0.9 |
| | 4 | Winter | | 100 | 55.0 ± 2.2 | 44.6 ± 2.1 | 0.4 ± 0.04 | 87.4 ± 4.9 |

| | | | | | | | | |
|---|---|---|---|---|---|---|---|---|
| *Acanthurus dussumieri* | 4 | Summer | 100 | | 4.4 ± 1.8 | 69.4 ± 2.4 | 26.2 ± 1.9 | 86.5 ± 1.7 |
| | 3 | Autumn | 100 | | 3.9 ± 2.0 | 60.3 ± 6.7 | 35.9 ± 6.4 | 92.4 ± 2.8 |
| | 3 | Winter | 100 | | 2.3 ± 0.7 | 73.7 ± 1.6 | 24.0 ±2.1 | 75.4 ± 12.1 |
| *Acanthurus lineatus* | 3 | Summer | 100 | | 1.6 ± 0.4 | 70.2 ± 0.5 | 28.2 ± 0.7 | 75.0 ± 0.9 |
| | 6 | Winter | 100 | | 1.5 ± 0.3 | 66.6 ± 4.3 | 32.4 ± 4.2 | 80.3 ± 2.5 |

**Supplementary Table 6:** Comparative relative opsin gene expression between winter and summer in seven surgeonfish (Acanthuridae) species: Expression levels (%) are normalised to two housekeeping genes: β-actin (*actb*) and elongation factor-1α (*elf1a*). Mean ± s.d.

| | | Cones | | | | | Rods |
|---|---|---|---|---|---|---|---|
| Species | Season | *sws2a* | *sws2b* | *rh2c* | *rh2a* | *lws* | *rh1* |
| *Acanthurus nigricauda* | Summer | 1.3±0.1 | | 0.1±0.03 | 8.4±1.5 | 4.4±0.6 | 59.4±2.8 |
| | Winter | 1.6±0.6 | | 0.3±0.1 | 10.8±4.0 | 5.0±5.0 | 66.3±11.7 |
| *Ctenochaetus striatus* | Summer | 1.0±0.2 | | 0.2±0.1 | 5.7±1.4 | 4.6±0.8 | 62.1±6.6 |
| | Winter | 1.6±0.3 | | 0.4±0.1 | 7.9±0.8 | 9.7±1.8 | 70.93.7± |
| *Naso brevirostris* | Summer | | 0.4±2.2 | 0.6±4.2 | 0.4±2.2 | 0.03±0.2 | 14.8±26.3 |
| | Winter | | 11.1±1.9 | 9.1±1.5 | 5.5±2.0 | 0.8±0.3 | 90.7±12.0 |
| *Zebrasoma veliferum* | Summer | 2.4±1.5 | | 0.4±0.4 | 5.7±3.6 | 6.8±3.9 | 63.9±21.0 |
| | Winter | 1.8±0.3 | | 0.2±0.1 | 8.4±2.2 | 5.7±1.8 | 78.4±13.5 |
| *Naso unicornis* | Summer | | 0.5±0.2 | 1.0±0.4 | 0.7±0.3 | 0.01±0.01 | 28.3±9.4 |
| | Winter | | 2.0±0.4 | 6.1±1.1 | 5.0±0.8 | 0.04±0.01 | 73.0±8.5 |
| *Acanthurus dussumieri* | Summer | 0.8±0.1 | | 0.4±0.1 | 6.4±1.5 | 2.3±0.3 | 69.0±5.1 |
| | Winter | 0.8±0.1 | | 0.2±0.1 | 5.7±0.5 | 1.9±0.1 | 48.3±20.4 |
| *Acanthurus lineatus* | Summer | 1.1±0.1 | | 0.3±0.1 | 10.6±0.3 | 4.3±0.2 | 51.8±2.3 |
| | Winter | 0.8±0.1 | | 0.2±0.01 | 7.0±1.1 | 2.9±0.5 | 42.1±5.5 |

**Supplementary Table 7:** Summary of annotations for the top 15 upregulated genes in summer compared with winter (i.e. largest fold change) for two species of surgeonfish: *Zebrasoma veliferum* and *Naso brevirostris*. Full gene functions (i.e. GO term) are provided (See Table S9).

| Species | Function | Gene symbol | logfc | Adj. p-val | Gene name |
|---|---|---|---|---|---|
| *Zebrasoma veliferum* | Complement component C1q complex binding | *ptx3* | 2.890 | 7.65E-03 | |
| | | *ptx3a* | 3.40 | 3.94E-03 | |
| | Membrane | *lrp12* | 3.20 | 2.42E-02 | Low density lipoprotein-related protein 12 |
| | DNA-binding transcription factor activity | *gata1a* | 3.34 | 3.21E-02 | GATA-type domain-containing protein |
| | Actin binding | *cald1a* | 3.36 | 1.51E-02 | Caldesmon 1a isoform X8 |
| | Oxygen carrier activity | *hbb1* | 3.47 | 3.36E-02 | Globin domain-containing protein |
| | Calcium ion binding | *anxa1a* | 4.51 | 5.46E-03 | Annexin |
| | | *syt5a* | 3.006 | 1.72E-02 | C2 domain-containing protein |
| | Uncharacterised | TRINITY_DN89599_c0_g1_i1 | 3.110 | 9.94E-03 | |
| | | TRINITY_DN59017_c0_g1_i2 | 3.43 | 3.94E-03 | |
| | | TRINITY_DN1586_c1_g1_i1 | 3.49 | 1.81E-02 | |
| | | TRINITY_DN279_c1_g1_i1 | 4.01 | 1.19E-02 | |
| | | TRINITY_DN1086_c0_g3_i1 | 4.03 | 3.58E-02 | |
| | | TRINITY_DN701_c0_g3_i1 | 3.367 | 3.94E-03 | |
| | | TRINITY_DN1710_c1_g2_i1 | 3.015 | 4.65E-02 | |
| *Naso brevirostris* | Oxygen carrier activity | *hba* | 5.17 | 3.19E-23 | Hemoglobin subunit alpha |
| | | *hbb1* | 4.31 | 5.32E-16 | Hemoglobin subunit beta-A |
| | | *hbb2* | 5.1 | 6.35E-16 | Hemoglobin subunit beta-B |
| | Calcium ion binding | *anxa5* | 4.64 | 2.59E-21 | Annexin |
| | | *rpl17* | 3.37 | 5.84E-27 | Large ribosomal subunit protein uL22 |

| Species | Function | Gene symbol | logfc | Adj. p-val | Gene name |
|---|---|---|---|---|---|
| *Naso brevirostris* | Structural constituent of ribosome/translation | *rpl9* | 2.79 | 2.99E-19 | Large ribosomal subunit protein uL6 |
| | | *rpl18* | 3.08 | 5.37E-17 | Large ribosomal subunit protein eL18 |
| | | *rpl6* | 2.63 | 3.73E-16 | Large ribosomal subunit protein eL6 |
| | | *rps11* | 2.74 | 9.65E-16 | Small ribosomal subunit protein uS17 |
| | | *rpl21* | 2.82 | 1.18E-15 | Large ribosomal subunit protein eL21 |
| | | *rplp2* | 2.71 | 2.07E-17 | Large ribosomal subunit protein P2 |
| | | *rpl11* | *2.57* | *2.35E-15* | Large ribosomal subunit protein uL11c |
| | RNA binding | *rpl19* | 3.05 | 2.07E-17 | Large ribosomal subunit protein eL19 |
| | | *rpl30* | 3.59 | 6.35E-16 | Large ribosomal subunit protein eL30 |
| | Mitochondrion | *rps3* | 2.54 | 9.22E-17 | Small ribosomal subunit protein uS3 |

**Supplementary Table 8:** Summary annotations for the top 15 downregulated genes in summer compared with winter (i.e. largest fold change) for two species of surgeonfish: *Zebrasoma veliferum* and *Naso brevirostris*. Full gene functions (i.e. GO terms) are provided (See Table S9).

| Species | Function | Gene symbol | logfc | Adj. p-val | Gene name |
|---|---|---|---|---|---|
| *Zebrasoma veliferum* | Polyubiquitin modification-dependent protein binding | *ubb* | -5.21 | 3.25E-06 | Polyubiquitin-B |
| | Protein binding | *ndrg1* | -3.66 | 1.63E-02 | Protein NDRG1 |
| | | *ndrg1-b* | -3.23 | 1.01E-02 | Protein NDRG1-B |
| | Acyltransferase activity | *lrata* | -3.65 | 9.63E-03 | Lecithin retinol acyltransferase a |
| | Transferase activity | *dtx4a* | -3.54 | 4.79E-04 | E3 ubiquitin-protein ligase |
| | Protein serine/threonine kinase activity | *dyrk4* | -3.34 | 1.33E-02 | Dual-specificity kinase |
| | Mitochondrion | *ndufa13* | -3.32 | 2.26E-02 | NADH dehydrogenase [ubiquinone] 1 alpha subcomplex subunit 13 |
| | | *si:dkey-283b15.2* | -3.03 | 1.72E-02 | Neuronal pentraxin-2 |
| | | *fam167b* | -2.97 | 2.08E-02 | Protein FAM167B |
| | Monooxygenase activity | *tph1* | -2.32 | 4.65E-02 | Us12 prolyl 3-hydroxylase |
| | Uncharacterised | TRINITY_DN104_c0_g1_i1 | -3.67 | 4.86E-03 | |
| | | TRINITY_DN1341_c0_g2_i4 | -3.47 | 3.94E-03 | |
| *Naso brevirostris* | Protein kinase activity | *camk1gb* | -6.79 | 5.37E-17 | Calcium/calmodulin-dependent protein kinase igb |
| | | *LOC127369744* | -4.97 | 1.92E-11 | Potassium voltage-gated channel subfamily B member 2-like |
| | SNAP receptor activity | *vamp2* | -3.98 | 3.49E-11 | Vesicle associated membrane protein 2 |
| | Calcium ion binding | *cdhr1* | -4.78 | 1.60E-09 | Cadherin-related family member 1 |
| | Transcription coregulator activity | *nrip1b* | -4.53 | 1.79E-09 | Nuclear receptor interacting protein |
| | | *LOC127371735* | -5.18 | 3.35E-09 | Dedicator of cytokinesis protein 3-like |
| | Metal ion binding | *slc6a1a* | -3.95 | 3.75E-09 | Solute carrier family 6 member 1a |
| | | *si:ch211-113d22.2* | -5.3 | 1.65E-25 | |
| | Protein binding | *ndrg1* | -4.4 | 0.037 | |

| Species | Function | Gene symbol | logfc | Adj. p-val | Gene name |
|---|---|---|---|---|---|
| *Naso brevirostris* | Uncharacterised | TRINITY_DN155_c0_g1_i1 | -3.97 | 9.92E-15 | |
| | | TRINITY_DN654_c0_g1_i4 | -6.05 | 2.21E-13 | |
| | | TRINITY_DN787_c0_g1_i1 | -4.09 | 2.95E-10 | |
| | | TRINITY_DN6859_c8_g1_i1 | -4.37 | 3.22E-10 | |
| | | TRINITY_DN581_c0_g2_i1 | -4.33 | 1.84E-09 | |
| | | TRINITY_DN126_c0_g1_i1 | -3.15 | 3.58E-09 | |
| | | TRINITY_DN545_c0_g1_i1 | -3.81 | 5.73E-09 | |

**Supplementary Table 9:** These were identified in the differential expression gene analysis (Table S7 and S8) of the surgeonfish species investigated.

| Go Term | Function |
|---|---|
| GO:0003779 | Actin binding |
| GO:0016746 | Acyltransferase activity |
| GO:0005509 | Calcium ion binding |
| GO:0001849 | Complement component C1q complex binding |
| GO:0000981 | DNA-binding transcription factor activity |
| GO:0016301 | Kinase activity |
| GO:0016020 | Membrane |
| GO:0046872 | Metal ion binding |
| GO:0005739 | Mitochondrion |
| GO:0004497 | Monooxygenase activity |
| GO:0005344 | Oxygen carrier activity |
| GO:0042461 | Photoreceptor cell development |
| GO:0005515 | Protein binding |
| GO:0031593 | Polyubiquitin modification-dependent protein binding |
| GO:0004672 | Protein kinase activity |
| GO:0004674 | Protein serine/threonine kinase activity |
| GO:0003723 | RNA binding |
| GO:0007165 | Signal transduction |
| GO:0005484 | SNAP receptor activity |
| GO:0003735 | Structural constituent of ribosome |
| GO:0003712 | Transcription coregulator activity |
| GO:0006412 | Translation |
| GO:001674 | Transferase activity |

Supplementary Methods

*Additional information on spectral sensitivity*

For surgeonfish RH1 opsins we identified three sites that were thought to cause $\lambda_{max}$ shifts compared to the primary reference sequence (*O. latipes* RH1) (D83N, A292S, S299A). All surgeonfish species showed D83N, which blue-shifts their RH1 pigments (-6 nm, Nathans 1990). Four species showed the substitution S299A, a site known to cause small blue-shifts (-2 nm, Fasick and Robinson 1998; Dungan et al. 2016). The short $\lambda_{max}$ of *C. binotatus* RH1 (484 nm) is due to A292S (-10 nm, Fasick and Robinson 1998; Janz and Farrens 2001).

We considered up to six sites that showed variations compared to the primary reference sequence (*O. niloticus* SWS2A) as potentially involved in SWS2A $\lambda_{max}$ tuning (M44T, I47V, I49V, M116T, L216F, S292A). M44T, a known tuning site in SWS2 (+4 nm, Yokoyama et al. 2007), red-shifted SWS2Aα in *Z. scopas*. All compared species showed I49V, a substitution known to blue-shift $\lambda_{max}$ in SWS2 (-2, Yokoyama and Tada 2003). Although not a known site, due to its proximity and similar nature we also considered I47V to cause a -2 nm shift in three species (*A. olivaceus*, *C. binotatus*, *A. pyroferus*). However, this site was only considered for the calculated lower boundary $\lambda_{max}$.

For surgeonfish SWS2B tuning we considered two sites that showed variations compared to the primary reference sequence (*O. niloticus* SWS2B), or among surgeonfish SWS2B sequences, as potentially causing $\lambda_{max}$ shifts (A109G, G164A). All investigated species showed these two substitutions, resulting in a net blue-shift of 1 nm compared to *O. niloticus* SWS2B (425 nm). Both, A109G (-2, Yokoyama et al. 2007) and G164A (+1, Yokoyama and Tada 2003) are known tuning sites in SWS2 opsins.

The effects of amino acid substitutions on spectral sensitivity tuning in RH2 visual pigments are not well understood (Yokoyama & Jia, 2020). Specifically, it seems that substitutions at many sites may cause $\lambda_{max}$ shifts when investigated in isolation, i.e., by site directed mutagenesis, but are either masked or cause entirely different spectral shifts when these substitutions occur in combination with other substitutions (e.g., Chinen et al. 2005; Yokoyama and Jia 2020). This is particularly relevant for inferred $\lambda_{max}$ values of RH2B visual pigments as these generally show more diversity than RH2A opsins (e.g., Stieb et al. 2016; Luehrmann et al. 2019). Here, we considered any known site effects that have been demonstrated via site directed mutagenesis, and hypothesized effects of substitutions at known tuning sites that showed unknown substitutions. Due to these, and as our approach does not account for possible combined site effects, the surgeonfish RH2B $\lambda_{max}$ values proposed here should be considered vastly hypothetical, and will require confirmation by different analytical means.

We considered 10 sites that showed variations compared to the lower limit reference sequence (*O. niloticus* RH2B) as potentially involved in $\lambda_{max}$ tuning of surgeonfish RH2B opsin (I49C, L60V, F74Y, M88C, I112V, A169C, V185C, L259M, T266V, S297A). Present in all species was I49C (-4 nm, Chinen et al. 2005). Also thought to cause blue-shifts were F74Y (-1 nm) in *P. hepatus*, as well as S297A (-2 nm) and T266V (-2 nm) in all species of the genus *Naso* (Chinen et al., 2005). Offsetting many of these effects in many species were several substitutions assumed to cause red-shifts. Chinen et al. (2005) demonstrated that the substitution T185C causes a 4 nm blue-shift in zebrafish RH2 opsins. All surgeonfish species assessed here showed 185C, however, *O. niloticus* (and *M. zebra*) RH2B had 185V. Considering that the substitution of an uncharged, polar/hydrophilic residue (T) for a moderately hydrophilic one (C) causes a small blue-shift (-4 nm; Chinen et al. 2005), we hypothesized that a substitution of a non-polar/hydrophobic residue for a moderately hydrophilic one (C) may in fact invert this effect, albeit at a smaller extent (+2 nm). Also thought to cause small red-shifts in a subset of the species investigated were: L60V (+2 nm, Chinen et al. 2005), present in all species except *N. tuberosus*, *N. annulatus*, *N. lituratus*, *A. triostegus* and *A. nigrofuscus*; M88C (+3 nm, Chinen et al. 2005), present in *A. blochii*, *A. dussumieri*, *A. xanthopterus*, *A. nigrocauda*, and *A. pyroferus*; I112V (+1 nm, Chinen et al. 2005), present only in *N. annulatus*, *N. tuberosus*, *N. caesius*, *N. unicornis*, *N. lituratus*, and *A. nigrofuscus*; A169C (+3 nm, Chinen et al. 2005), present only in several Naso species (*N. caesiusi*, *N. unicornis*, *N. vlamingii*, *N. brevirostris*, *N. hexacanthus* and *N. lituratus*. Chinen et al. (2005) demonstrated that substituting V for A at site 169 causes small (-1 nm) blue-shifts. We expect that the substitution of a polar amino acid (C) causes a stronger and opposite effect. L259M, found only in *A. lineatus*, *C. binotatus*, *P. hepatus*, *Z. scopas*, and *Z. veliferumum*, is one of nine amino acid changes that combined are known to cause a strong red-shift (+20 nm) in RH2 opsins (Yokoyama & Jia, 2020). As one of these (Q122E) is known to cause a +15 nm shift on its own (Yokoyama & Jia, 2020), we estimated L259M to cause a 1 nm red-shift.

The upper limit reference sequence (*M. zebra* RH2B) showed three additional (I44M, S46F, I205M) and two different (I88C, F259M) variations that were considered for surgeonfish $\lambda_{max}$ tuning. I44M, seen in all surgeonfish species assessed here, was thought to possibly blue-shift $\lambda_{max}$ by 1 nm, as this is an inversion of a substitution known to be involved in a 3 nm red-shift facilitated by a complex of substitutions at other sites (Yokoyama & Jia, 2020). Part of the same site effect complex, I205M, also present in all surgeonfish species, is thought to offset the shift caused by I44M (+1 nm, Yokoyama and Jia 2020). Compared to *M. zebra* RH2B all surgeonfish species showed S46F, which was estimated to act in a similar way as the known RH2 substitution L46F (-3 nm, Chinen et al. 2005). *M. zebra* RH2B had Isoleucin at site 88, however, I88C was estimated to be similar in effect as M88C (see above). F259M was thought to have a similar effect as L259M (+1 nm, see above).

For surgeonfish RH2A tuning we considered up to six sites (two for upper, six for lower limit) that showed variations compared to the primary reference sequence (*O. niloticus* RH2Aβ) as possibly causing $\lambda_{max}$ shifts (F60L, C88A, V209C, M259F/C, T266V, G273A). To calculate the upper limit estimate, we considered only two sites which had previously been investigated via site directed mutagenesis to identify single site substitution effects (C88A, T266V). The additional four sites used to calculate the lower spectral sensitivity limits have also been shown to be involved in RH2 tuning in teleosts (Yokoyama & Jia 2020). However, their effects have only been described in conjunction (+3 nm) whereas the individual effects of these sites are not known. Surgeonfish show substitutions that constitute an inversion of polarity compared to the substitutions described by Yokoyama and Jia (2020). Considering the overall small shift attributed to all of these sites acting together (+3 nm), we tentatively assumed that each of these may slightly blue-shift (-1 nm) $\lambda_{max}$ of the respective surgeonfish RH2A pigments.

We considered two sites that showed variations compared to the primary reference sequence (*O. niloticus* LWS) as potentially involved in $\lambda_{max}$ tuning (S164A, T269A). *A. pyroferus* was the only species showing S164A, a known LWS tuning site (-7 nm, Asenjo, Rim, & Oprian, 1994). The short $\lambda_{max}$ estimated for *N. lituratus*, *N. unicornis* and *N. vlamingii* were due to T269A, a known LWS tuning site substitution that strongly blue-shifts $\lambda_{max}$ (-16 nm, Asenjo et al., 1994).
